# Myf6/MRF4 is a Myogenic Niche Regulator Required for the Maintenance of the Muscle Stem Cell Pool

**DOI:** 10.1101/691386

**Authors:** Felicia Lazure, Darren M. Blackburn, Nabila Karam, Korin Sahinyan, Ahmad Sharanek, Duy Nguyen, Aldo H. Corchado, Christoph Lepper, Hamed S. Najafabadi, Theodore J. Perkins, Arezu Jahani-Asl, Vahab Soleimani

## Abstract

In metazoans, skeletal muscle evolved to contract and produce force. However, recent experimental evidence suggests that skeletal muscle has also acquired endocrine functions and produces a vast array of myokines. Using ChIP-Seq and gene expression analyses of myogenic factors, we show that Myf6/MRF4 transcriptionally regulates a broad spectrum of myokines and muscle-secreted proteins, including ligands for downstream activation of key signaling pathways such as EGFR, STAT3 and VEGFR. Homozygous deletion of Myf6 causes a significant reduction in the ability of muscle to produce key myokines such as EGF, VEGFA and LIF. Consequently, although Myf6 knockout mice are born with a normal muscle stem cell compartment, they undergo progressive reduction in their stem cell pool during postnatal life. Mechanistically, muscle stem cells from the Myf6 knockout animals show defects in activation of EGFR and STAT3 signaling, upregulate the p38 MAP kinase pathway and spontaneously break from quiescence. Exogenous application of recombinant EGF and LIF rescue the defects in the muscle stem cell pool of Myf6 knockout animals. Finally, skeletal muscles of mice lacking Myf6 have a significantly reduced ability to sustain donor-engrafted muscle stem cells. Taken together, our data uncovers a novel role for Myf6 in regulating the expression of niche factors and myokines to maintain the skeletal muscle stem cell pool in adult mice.

## Introduction

In adult skeletal muscle, Muscle stem cells (MuSCs) are spatially housed between the sarcolemma of the muscle fibers and the basal lamina, a layer of extracellular matrix proteins (ECM) that are closely juxtaposed with the myofibers. The physical enclosure of MuSCs in such an anatomical space ensures that they remain in close proximity to the myofiber in a mitotically quiescent state, under normal physiological conditions. Injury or trauma to the myofiber causes disruption of the MuSC niche during which they re-enter cell cycle, transiently proliferate and undergo myogenic differentiation to repair the injured muscle ^1-3^. Induction of MuSC activation triggers a temporally regulated myogenic differentiation program orchestrated by four myogenic factors, namely Myf5, MyoD, Myogenin and Myf6 ^4-6^. MyoD and Myf5 transiently amplify MuSCs but also induce differentiation by upregulating Myogenin in a context dependent manner ^7,8^. While Myf5, MyoD and Myogenin are transiently expressed during myogenic commitment and differentiation of MuSCs, in the fully differentiated myofiber, Myf6/MRF4 is the predominant myogenic factor that remains expressed under physiological conditions. Previous studies have implicated Myf6 involvement in myofiber maturation and in the maintenance of their differentiated state ^9^. Aside from its function in differentiation, Myf6 has also been implicated as a muscle determination factor in the absence of MyoD and Myf5 ^10^.

While the epigenetic and transcriptional control of muscle formation has been extensively studied ^11,12^, less is known about the precise mechanisms by which the muscle stem cell pool is maintained in adult skeletal muscle. Maintenance of the MuSC pool crucially depends on a dynamic balance between quiescence and activation ^13^, a process that is intricately dependent on the interaction of MuSCs with their niche microenvironment ^14-16^. Many non-myogenic cells have been shown to contribute to the niche milieu of MuSCs, such as endothelial cells under normal physiological conditions ^15^, fibroblasts under hypertrophy ^17^ and macrophages and neutrophils during muscle injury ^18-20^. The physical enclosure of MuSCs by the ECM creates an anatomically defined niche microenvironment where the muscle stem cell resides in a quiescence state. However, how molecular communication between myofibers and their associated MuSCs regulates their fate remains unknown.

Beyond its contractile properties, skeletal muscle plays an important role as an endocrine organ. The secretome of skeletal muscle is composed of proteins and peptides known as myokines, which are thought to have system-wide effects through autocrine, paracrine, and endocrine functions ^21,22^. Exercise-induced myokines such as Interleukin-6 (IL6), Interleukin-15 (IL15), and brain-derived neurotrophic factor (BDNF) among others, have been studied in the context of their effect on inflammation, metabolic homeostasis and lipid oxidation, respectively ^23-25^.

Here, we show that Myf6 plays a key role in the endocrine function of skeletal muscle by transcriptionally regulating multiple myokines such as EGF, LIF and IL6. Our data shows that regulation of myokine expression by Myf6 is required to maintain the adult muscle stem cell pool. In the absence of Myf6, MuSCs break quiescence, upregulate the p38 MAPK signaling pathway and undergo differentiation. Rescue of MuSC numbers by *in vivo* exposure to exogenous myokines in acute muscle injury experiments suggests that defects in the MuSCs compartment in Myf6 knockout animals are principally due to defective myokine signaling. Collectively, our data suggest that Myf6 has a novel function as a myogenic niche regulator and plays a critical role in regulating stem cell pool in adult skeletal muscle.

## Results

### Myf6/MRF4 Cistrome and Associated Genetic Networks in Skeletal Muscle

Among the four myogenic regulatory factors (MRFs), Myf6 is the only factor that is primarily expressed in fully differentiated muscle fibers (Fig. 1A). The remaining three MRFs, namely Myf5, MyoD and Myogenin are transiently expressed during muscle stem cell activation, commitment and terminal differentiation. To identify genome-wide Myf6 targets in primary myotubes, we first used Chromatin Tandem Affinity Purification Sequencing (ChTAP-Seq), an ideal alternative to traditional ChIP-Seq when a commercial ChIP-grade antibody is not available. Here, we generated primary myoblast cell lines over-expressing a C-terminal Tandem Affinity Purification tag (Myf6-CTAP), using a retroviral vector. We validated the expression, subcellular localization and functionality of the ectopic Myf6-CTAP in muscle and non-muscle cells using immunofluorescence (IF), Western Blotting (WB) and Luciferase assays (Fig. S1). Next, we performed ChTAP-Seq on multinucleated primary myotubes formed after five days in differentiation media as described previously ^26^. Using MACS2 ^27^ with a p-value threshold of 10^−5^ and default parameters, we identified 12,885 binding sites for Myf6 in primary myotubes, using sequencing reads in the affinity-matched empty vector control (EV-ChTAP-Seq) as background (Fig. 1A). Motif analysis of the peak-covered sequences from the total number of Myf6 peaks using MEME suite ^28^ identified the top scoring canonical E-box motif, centrally located and closely juxtaposed by the MEF2A binding motif located peripheral to the E-box counterparts (Fig. 1B). Comparative analysis between the densities of the E-box motif distribution in the Myf6 ChIP-Seq peaks compared to random genomic blocks showed significantly denser clustering of E-boxes under Myf6 peaks (Fig. 1C). Such patterns of motif distribution in Myf6 peaks mirror those of other myogenic factors such as MyoD and Myf5 ^6^. Within Myf6 peaks, highly enriched motifs for known MRF cooperative heterodimers such as E-protein (E12/E47) and antagonistic interactors such as TWIST1 and ID4 followed similar distribution patterns as those of E-box motifs (Fig. 1D). Together, these data suggest that transcriptional control of gene expression by Myf6 follows similar operational mechanisms as those deployed by other MRFs within their cis-regulatory domains.

**Figure 1.**
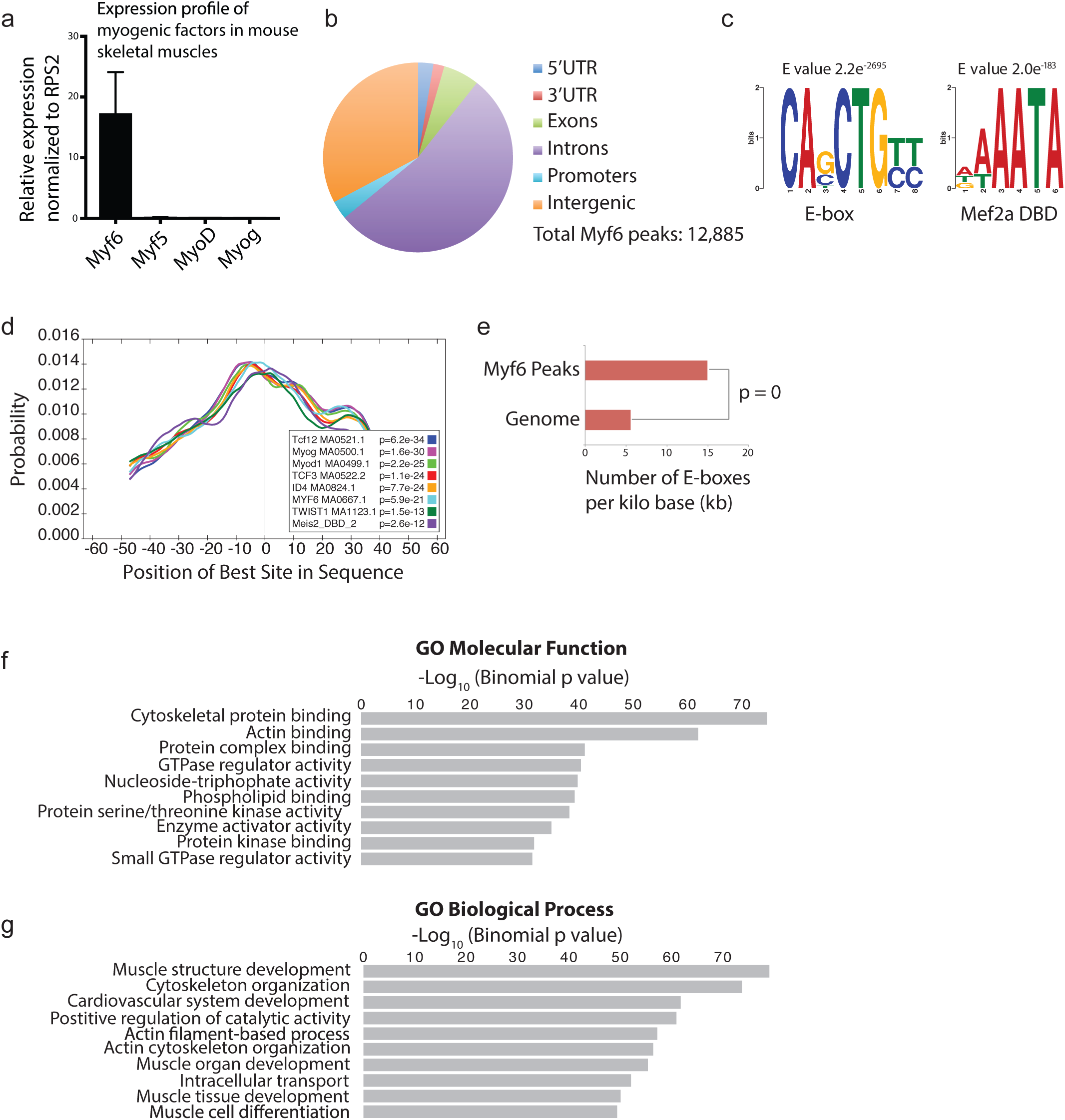
Genome-Wide Binding Sites of Myf6/MRF4 in Primary Myotubes. **(a)** Relative expression of myogenic factors in mouse hindlimb skeletal muscle assayed by quantitative real-time PCR (RT-qPCR) on isolated whole muscle RNA. **(b)** A pie chart showing the distribution of Myf6 ChIP-Seq peaks within various genomic features. **(c)** Motif analysis using MEME suite ^28^ identified canonical E-box motif as the most highly enriched sequence among all sequences under Myf6 peaks, followed by the Mef2a DNA binding motif. **(d)** Spatial distribution of transcription factor binding motifs within Myf6 ChIP-Seq peaks using CentriMo available via MEME suite ^28^. **(e)** Normalized distribution of E-box motifs that occur within Myf6 peaks compared to their occurrences in the genome normalized per kilo base of DNA. **(f)** Gene Ontology (GO Molecular Function) analysis of genes associated with Myf6 peaks based on association by proximity (single gene within 100kb of peaks) analyzed by Genomic Regions Enrichment of Annotations Tool (GREAT) ^29^. **(g)** Gene Ontology (GO Biological Process) analysis of genes associated with Myf6 peaks based on proximity similar to the analysis in “f”.

### Myf6 Occupies Regulatory Domains of Functionally Distinct Sets of Genes in Differentiated Muscle Cells

To identify the genetic networks regulated by Myf6 in differentiated muscle cells, we first used Genomic Regions Enrichment of Annotations Tool (GREAT) ^29^ and performed Gene Ontology (GO) analysis using an association rule of single nearest genes (300kb maximum extension). Within those regions, 12,270 out of 12,885 peaks (95.2%) were associated with a single gene and 610 peaks (4.73%) were not associated with any gene. Computation of GO terms using binomial tests, implemented in GREAT ^29^ showed that the majority of the Myf6 ChIP-Seq peaks in the primary differentiated myotubes were within the regulatory domains of genes involved in regulation of muscle structure, muscle cell development and differentiation, as well as actin filament organization, as expected (Fig. 1f, 1g). However, in addition to the expected differentiation genes, we identified a large set of genes for secreted proteins including many known myokines and chemokines that are bound and transcriptionally regulated by Myf6 (Fig. 2a, Fig. S2). The latter genes code for ligands such as EGF, LIF, and IL6 among others, which activate EGFR and STAT3 signaling pathways. Both EGFR and STAT3 play important roles in the regulation of muscle stem cell self-renewal and stem cell pool^30,31^. This data suggests that a novel function of Myf6 in adult skeletal muscle may be the establishment of a myokine-mediated regulatory network, which signals back to MuSCs. Therefore, we hypothesize that genetic deletion of Myf6 in skeletal muscle leads to impairment in EGFR and STAT3 signaling in MuSCs due to diminished expression of myokines. Other myokines such as Vascular endothelial growth factor (VEGFA) are also transcriptionally regulated by Myf6 (Fig. 2a, 2b, 2f, S2a-e). VERGFA plays an important role in maintaining MuSCs quiescence via recruitment of endothelial cells^15^.

**Figure 2.**
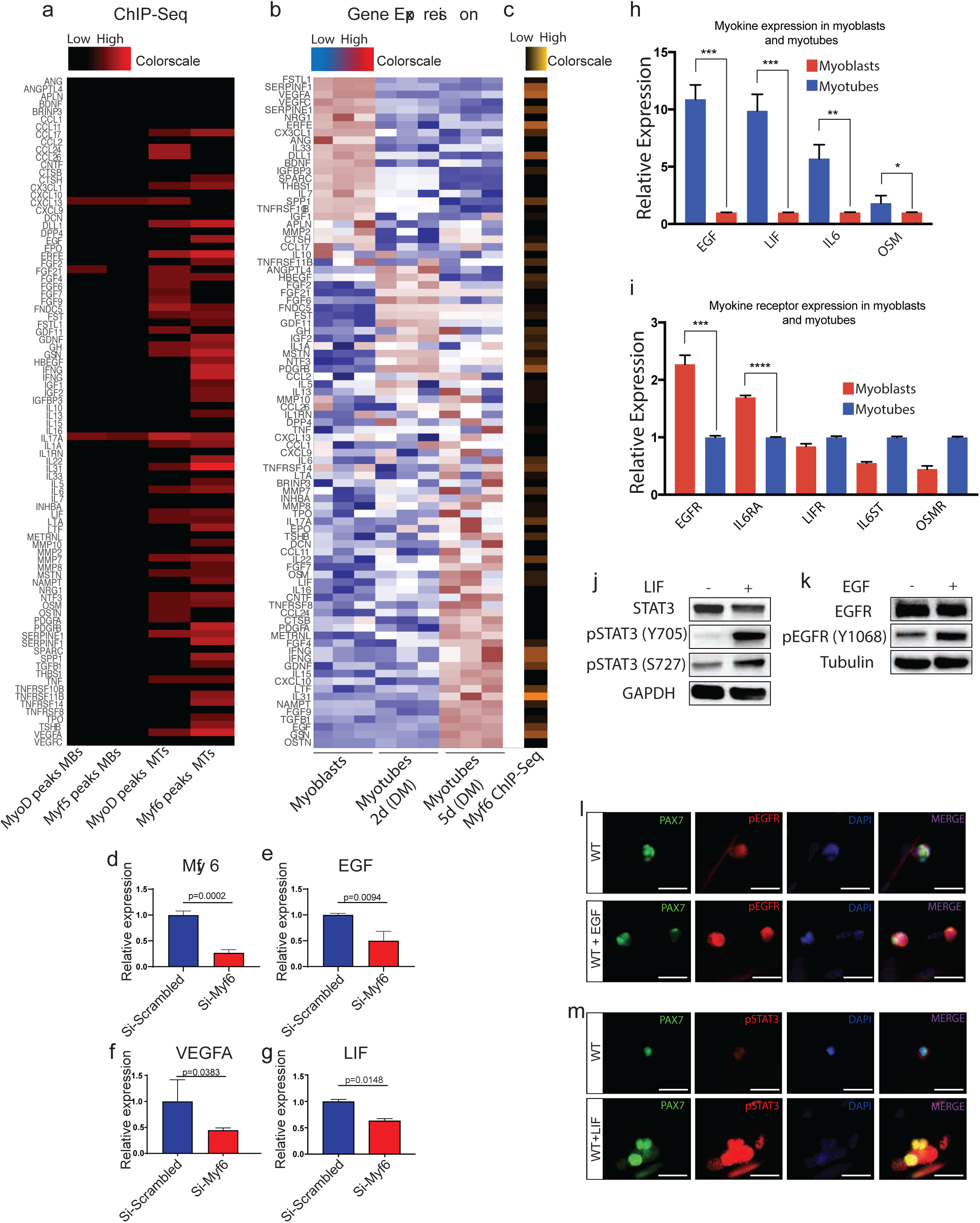
Temporally Regulated Occupancy of Myogenic Regulatory Factors (MRFs) to cis-Regulatory Domains of Myokine Genes. **(a)** Colormap of Myf5, MyoD and Myf6 peaks within 100kb of the Transcription Start Sites (TSS) of myokines (supplemental table S2) ranging from zero (black) to six peaks (red) occupancy. The onset of differentiation coincides with increased binding of MRFs to the regulatory domains of the myokines genes. **(b)** Gene expression analysis of myokines (Supplemental table S2) during a five day time course of myogenic differentiation going from cycling myoblasts in growth media (Ham’s F10 supplemented with 20% Fetal Bovine Serum, 1% penicillin/streptomycin, 2.5 ng/ml basic Fibroblast Growth Factor) to terminally differentiated myocytes (2 days in differentiation media, DMEM supplemented with 5% horse serum) to post mitotic multinucleated myotubes (5 days in differentiation media). Gene expression was assayed in biological triplicate by microarray ^7,36^. Expression of each gene was averaged across three replicates and normalized to mean zero and standard deviation of one. Expression values were then truncated at +/− 3 for color display. Blue indicates lower than average expression, while red indicates higher than average expression. **(c)** Distribution of Myf6 peaks within 100kb of the TSS of the myokine genes (supplemental table S2), similar to the analysis shown in “a”. **(d)** Depletion of Myf6 transcript by siRNAs in differentiated primary myotubes (5DM). **(e-g)** Depletion of Myf6 expression by siRNAs leads to a significant reduction in the gene expression output of Epidermal Growth Factor (EGF) **(e)**, Vascular Endothelial Growth Factor A (VEGFA) **(f)** and Leukemia Inhibitory Factor (LIF) **(g). (h)** Quantitative Real-Time PCR (RT-qPCR) analysis of a select set of myokines genes involved in the activation of STAT3 and EGFR signaling pathways. RT-qPCR analysis was performed on the total RNA isolated from primary myoblasts and multinucleated primary myotubes derived from MuSCs isolated by Fluorescence Activated Cell Sorting (FACS, see the Extended Material and Methods). Myotubes were obtained by feeding >90% confluent primary myoblasts with differentiation media (DMEM supplemented with 5% horse serum) for five days. **(i)** RT-qPCR analysis of the expression of select receptors for the ligands in “h” between primary myoblasts and multinucleated myotubes. **(j-k)** Activation of STAT3 and EGFR signaling in myoblasts treated with recombinant LIF **(j)** and EGF **(k)** respectively. Myoblasts were treated with 2 ng/ml of each recombinant protein in growth media (Ham’s F10 supplemented with 20% FBS, 2.5 ng/ml bFGF) for 15 minutes and lysed in RIPA lysis buffer. Western Blots were performed with antibodies against EGFR, phospho-EGFR (pEGFR-Y1068), STAT3 and phospho-STAT3 (pSTAT3-Y705). **(l)** Activation of EGFR in MuSCs on EDL myofibers treated with EGF compared to untreated control. Myofibers were cultured for 48 hours and were treated with 40ng/ml of EGF for 10 minutes before fixing and staining with antibodies for PAX7, EGFR and phospho-EGFR. **(m)** Activation of STAT3 signaling in MuSCs on EDL myofibers treated with LIF compared to untreated control. Myofibers were cultured for 48 hours and were treated with 1000U/ml of recombinant LIF for 10 minutes before fixation and staining with antibodies for PAX7, STAT3 and p-STAT3.

*In vivo* depletion of Myf6 transcript by RNAi in differentiating primary myotubes shows that Myf6 is required for transcriptional regulation of VEGFA, EGF and LIF (Fig. 2d-g). While some ligands such as VEGFA are produced by both progenitors as well as differentiated myotubes (Fig. 2b, 2f, S2b), others such as EGF and LIF are principally produced in differentiated myotubes (Fig. 2b, 2h). Interestingly, while EGF, LIF, OSM and IL6 are expressed in differentiated myotubes, their corresponding receptors are expressed in progenitors (Fig. 2i). This observation that differentiation of muscle stem cells creates a physical niche whereby myokines (ligands) are produced in myofibers while their respective receptors are expressed in their associated MuSCs suggests the existence of a myokine-mediated communication network between myofiber and MuSCs. Below, we delve into the critical role of Myf6 in establishing a myokine signaling network which is required for activation of EGFR and STAT3 signaling pathways in MuSCs.

### Myf6-Regulated Myokines Activate EGFR and STAT3 Signaling Pathways in Muscle Stem Cells

EGFR and STAT3 have recently been shown to play crucial roles in regulating muscle stem cell self-renewal and expansion. Both EGFR and STAT3 are readily activated in proliferating myoblasts (Fig. 2j) as well as in myofiber-associated MuSCs (Fig. 2i, 2m) in response to exogenously applied ligands, indicating that these pathways are operationally active in muscle cells. To determine the requirement of Myf6 in the regulation of myokine expression *in vivo* we first analyzed the whole muscle transcriptome of Myf6-knockout mice under normal physiological conditions and after cardiotoxin (CTX) injury by RNA-Seq. For this, we used Myf6^CE^ mice, in which a Cre-ER^T2^ cassette is knocked into the first exon of the *Myf6* gene, rendering it a null allele (Fig. 3a)^32^. Mice with biallelic deletion of Myf6 were born with normal skeletal muscles and were indistinguishable from their WT littermates (Fig. 2b). Quantitative Real-Time PCR (RT-qPCR) of RNA extracted from hindlimb muscles showed efficient depletion of Myf6 transcript in homozygous Myf6 knockout mice (Fig. 3c). RNA-Seq analysis showed that genetic deletion of Myf6 in skeletal muscles led to deregulation of a vast array of genes involved in myoblast differentiation, muscle development, muscle contraction, and pre-mRNA splicing (Table S3). However, our data suggests that a novel function of Myf6 is to regulate the expression of myokines such as VEGFA, EGF, LIF, IL-15 and IL-6 (Fig. 2a, 2b, 2d-g, 3d, S2a-d). Analysis of Myf6 binding sites by ChIP-Seq showed enrichment of Myf6 peaks around the transcription start sites (TSS) of latter genes (Fig. 3f). Taken together, this data suggests that, in addition to regulating muscle differentiation genes, Myf6 transcriptionally regulates myokines that play key roles in the activation of signaling pathways such as EGFR and STAT3. Using both siRNA to specifically knockdown Myf6 in myotubes (Fig. 2d-g) and *in vivo* genetic deletion of Myf6 in skeletal muscle followed by whole muscle RNA-Seq analysis (Fig. 3d, S2), we show that the unique function of Myf6 in regulating myokine expression is not compensated by other myogenic factors such as MyoD and Myogenin. Next, to determine if loss of Myf6 leads to diminished myokine protein production, we isolated MuSCs from Myf6-KO and WT counterparts by Fluorescent Activated Cell Sorting (FACS) using ITGA7^+^/Lin^−^ as described in Materials and Methods. We then differentiated fully confluent cultures of primary myoblasts isolated from Myf6-KO and WT animals for seven days in differentiation media (DMEM supplemented with 5% horse serum). Next, we collected media (secretome) from myotubes and performed Enzyme-Linked Immunosorbent Assay (ELISA) followed by comparative analysis of protein expression by normalizing protein amount to the number of nuclei as described in Materials and Methods. ELISA analysis of two key myokines, EGF and VEGFA shows that the Myf6-KO secretome has a significant lower level of these myokines compared to their WT counterparts (Fig. 3f, 3g). Similarly, to determine the overall contribution of Myf6 to the level of myokines in serum, we isolated blood serum from Myf6-KO and WT counterpart and by ELISA analysis showed that serum of Myf6-KO mice has a significant reduction in key myokines such as EGF and VEGFA (Fig. 3h, 3i). This data suggests that Myf6 is required for the production of multiples myokines.

**Figure 3.**
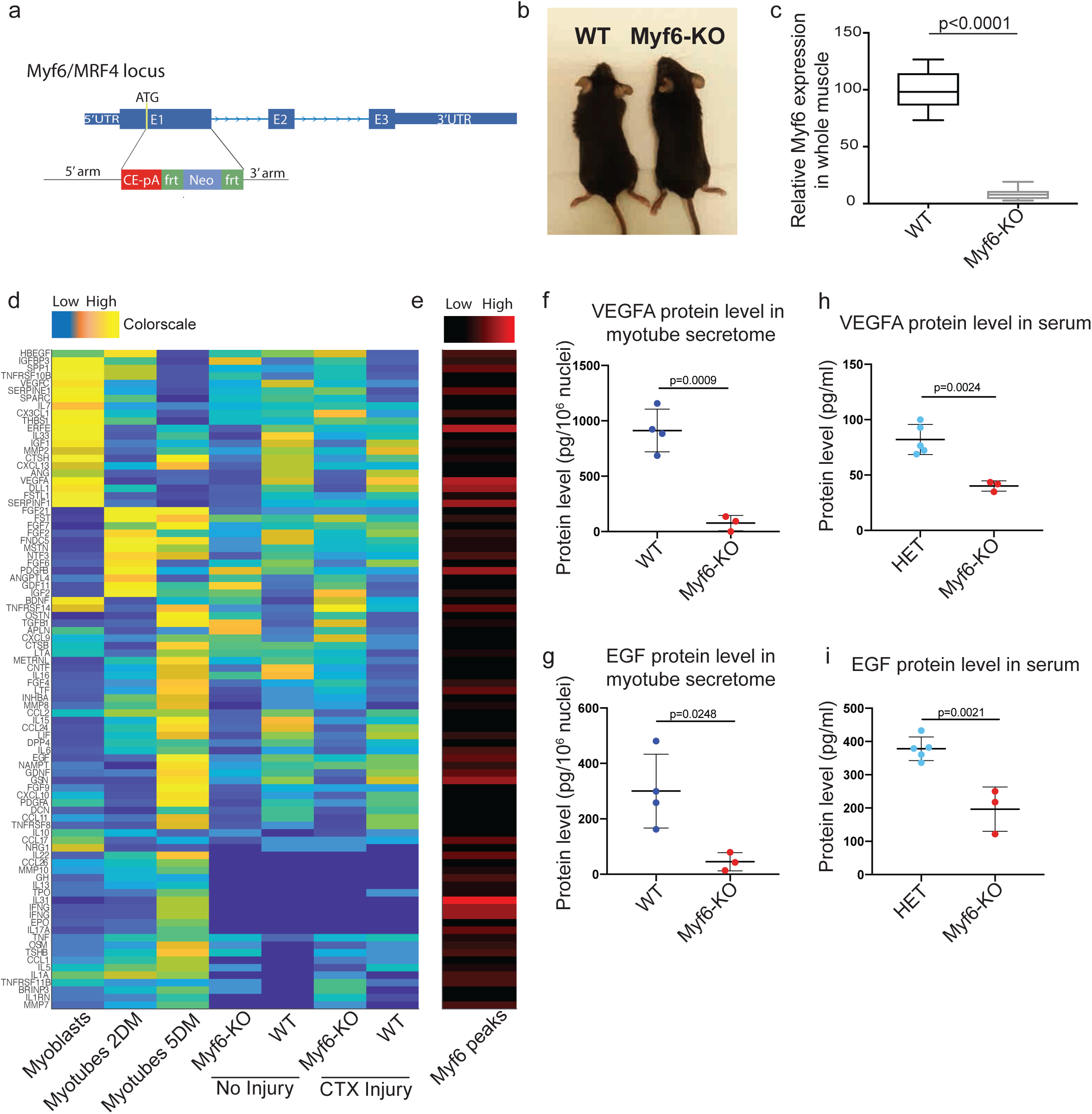
Genetic Deletion of Myf6 Impairs Myokine Expression in Adult Skeletal Muscles. **(a)** Schematic drawing of the Myf6 locus showing insertion of Cre-ER^T2^ sequence rendering the Myf6 locus as a null allele ^32^. **(b)** Picture of 8 week-old Myf6-KO and WT mice. **(c)** Relative expression of Myf6 transcript in the hindlimb skeletal muscles of Myf6-KO and WT mice measured by Quantitative Real-Time PCR (RT-qPCR). **(d)** Color map of genes that are up/down regulated in the whole muscle transcriptome (RNA-Seq) of hindlimb muscles from Myf6-KO and WT counterparts. Differential expression is Log_2_ transformed and truncated at +/−3. Color scale goes from blue (−3=lower than mean), to white (0=mean) and finally to red (+3=higher than mean). **(e)** Distribution of Myf6 peaks within 100kb of the TSS of the myokine genes ranging from zero (black) to six ChIP-Seq peaks (red) occupancy. **(e-f)** Expression of VEGFA **(e)** and EGF **(f)** proteins in the secretome of Myf6-KO and WT myotubes as measured by enzyme-linked immunosorbent assay (ELISA). Primary myotubes were derived from MuSCs isolated by Fluorescence Activated Cell Sorting (FACS, see the Extended Material and Methods) from Myf6-KO and WT mice. Myotubes were obtained by feeding >90% confluent primary myoblasts with differentiation media (DMEM supplemented with 5% horse serum) for seven days. Media from the last 48 hours of differentiation (days 5 to 7) was collected for ELISA secretome analysis. Protein readouts were normalized to the total number of nuclei in each plate. **(h-i)** Expression of VEGFA **(h)** and EGF **(i)** proteins in the blood serum from Myf6-KO and heterozygous counterparts as measured by Enzyme-Linked Immunosorbent Assay (ELISA).

### Loss of Myf6 Leads to Impaired Activation of EGFR and STAT3 Signaling Pathways in the Muscle Stem Cells

Our data thus far suggests that Myf6 is indispensable for the expression of multiple myokines in skeletal muscles. These myokines such as EGF and LIF regulate critical signaling pathways such as EGFR and STAT3. Thus, we hypothesized that loss of Myf6 can lead to impairment in activation of EGFR and STAT3 signaling pathways in myofiber-associated MuSCs. To determine the role of Myf6 in transcriptional regulation of myokines, we first analyzed Myf6 binding sites in their vicinity (Fig. 4a, 4b). Notably, our ChIP-Seq data shows strong enrichment of Myf6 ChIP-Seq reads, but not the reads from the affinity matched control ChIP-Seq, on the enhancer elements in the vicinity of the TSS of EGF, LIF, OSM and VEGFA in primary myotubes (Fig. 4a-b). Next, we determined the pattern of regulatory histone marks including Histone H3 mono methyl lysine 4 (H3K4me1), a marker for enhancer elements and histone H3 trimethyl lysine 4 histone (H3K4me3), marking active/poised TSS (Fig. 4a-c). Notably, our analysis of ChIP-Seq data indicates that Myf6 binding sites overlap with H3K4me1 (Fig. 4a-c). Taken together, the analysis of ChIP-Seq data (Fig. 4a-c), gene expression data by loss of function of Myf6 (Fig. 2) and analysis of protein expression by ELISA (Fig. 3f-i) shows that Myf6 is required for the expression of key myokines. Given that ligands for activation of EGFR and STAT3 are under transcriptional control of Myf6 in differentiated muscle cells (Fig. 2, Fig. 3, Fig. 4a-c) we hypothesized that genetic deletion of Myf6 in the skeletal muscle myofiber can lead to defects in phosphorylation and subsequent activation of EGFR and STAT3. To test this hypothesis, we isolated EDL myofibers from Myf6-KO and WT animals and cultured them in growth media for 44 hours. To eliminate the compounding effect of mitogens and growth factors in fiber culture, we switched fibers to pure DMEM for four hours before fixing and staining. Immunofluorescence of EDL myofibers shows defects in phosphorylation of EGFR (pEGFR-Y1068) and STAT3 (pSTAT3-Y705) (Fig. 4d-g). Importantly, treatment of EDL myofibers from Myf6-KO animals with recombinant EGF or LIF rescues defects in EGFR and STAT3 phosphorylation, respectively (Fig. 4e-g). This data suggests that Myf6 is required to maintain functional EGFR and STAT3 in the myofiber-associated MuSCs.

**Figure 4.**
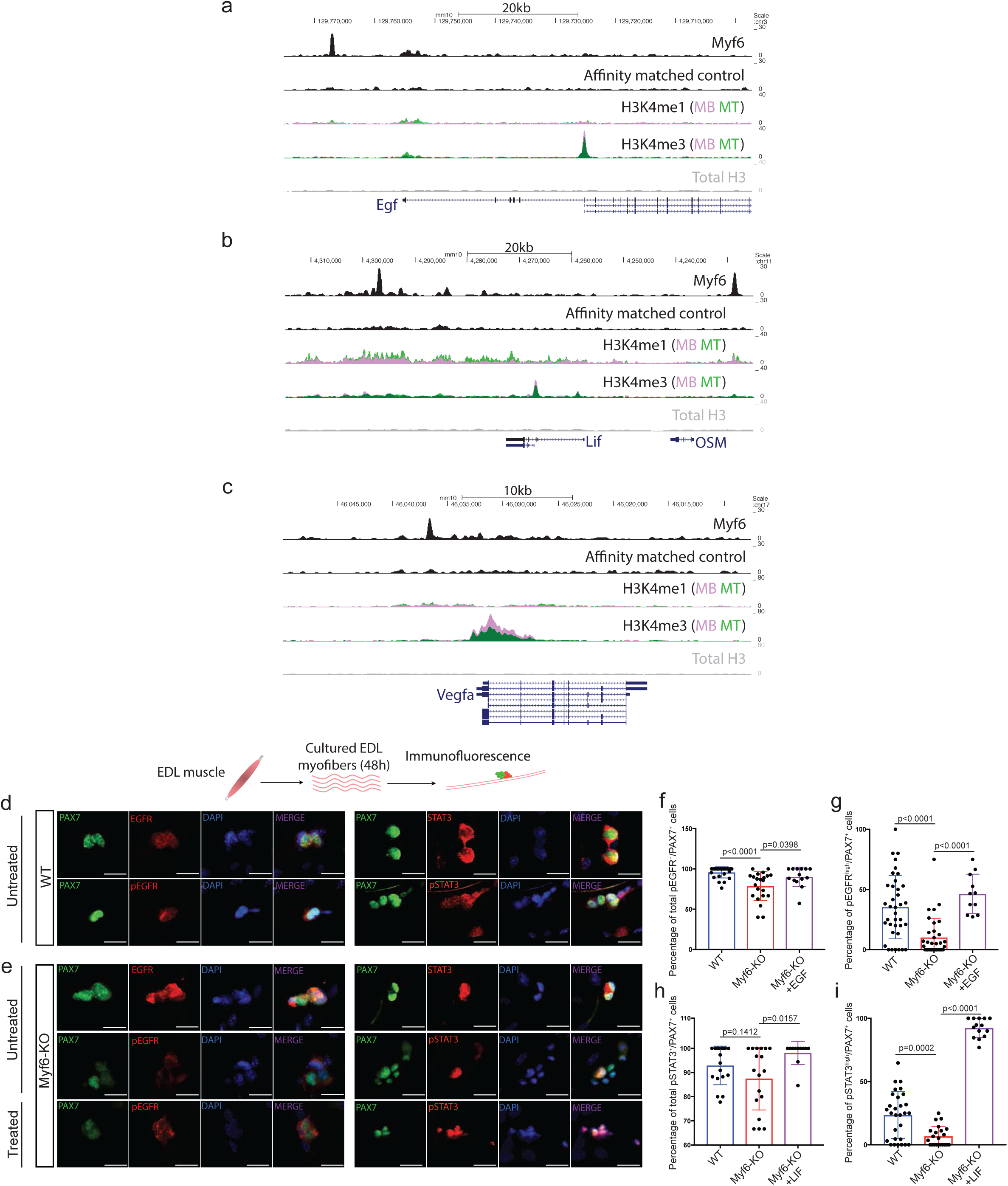
Myf6 Activates Ligands are Required for Regulation of Key Canonical Signaling Pathways in MuSCs. **(a-c)** Snapshots of the UCSC genome browser showing enrichment of Myf6 ChIP-Seq reads compared to control around EGF **(a)**, LIF and OSM **(b)**, and VEGFA **(c)** loci superimposed on reads for Histone H3 lysine 4 mono methyl (H3K4me1), Histone H3 lysine4 tri methyl (H3K4me3) and total H3 in myoblasts (pink) and myotubes (green). **(d-e)** Immunofluorescence staining of WT **(d)** and Myf6-KO **(e)** EDL myofibers stained with PAX7, EGFR, phospho-EGFR, STAT3 and phospho-STAT3 antibodies. Live fibers were maintained in culture media for 44 hours, followed by 4 hours of serum starving in non-supplemented DMEM. For treated samples, 40ng/ml of recombinant EGF (left) or 1000U/ml of recombinant LIF (right) was added to the non-supplemented DMEM for 10 minutes before fixation. Activation status of EGFR and STAT3 signaling pathways was assessed using antibodies against phospho-EGFR (pEGFR-Y1068) and phospho-STAT3 (pSTAT3-Y705), respectively. **(f)** Quantification of the percentage of all pEGFR^+^/PAX7^+^ cells (defined as PAX7^+^ cells showing low or high pEGFR signal) on WT, Myf6-KO, and Myf6-KO EDL myofibers treated with EGF. **(g)** Quantification of the percentage of EGFR^high^/PAX7^+^ cells (defined as PAX7^+^ cells showing high pEGFR signal) on WT, Myf6-KO, and Myf6-KO EDL myofibers treated with EGF. **(h)** Quantification of the percentage of all pSTAT3^+^/PAX7^+^ (defined as PAX7^+^ cells showing low or high pSTAT3 signal) cells on WT, Myf6-KO, and Myf6-KO EDL myofibers treated with LIF. **(i)** Quantification of the percentage of pSTAT3^high^/PAX7^+^ cells (defined as PAX7^+^ cells showing high pSTAT3 signal) on WT, Myf6-KO, and Myf6-KO EDL myofibers treated with LIF.

### Genetic Deletion of Myf6 Leads to Exhaustion of the MuSC Pool in Adult Skeletal Muscle

Genetic deletion of Myf6 led to down-regulation of many myokine genes including *EGF, HBEGF, LIF, VEGFA* and *Il-15* (Fig. 2, Fig. 3). Therefore, we analyzed how the loss of Myf6 affects muscle regeneration and the stem cell pool. Mice with a homozygous deletion of Myf6 were born without any visible abnormalities and were fertile (Fig. 3). Immunofluorescent analysis of cross sections of TA muscles showed no significant difference in the number of regenerating fibers (i.e., myofibers with centrally located myonuclei), or the number of myofibers per unit area (mm^2^) between Myf6-KO and WT mice (Fig. S3). These data suggest that Myf6 is dispensable for muscle formation and MuSC differentiation during development. Next, we analyzed the effect of genetic deletion of *Myf6* on the MuSC pool postnatally (P7, P21) and during adulthood (three months and 10 months of age) by first performing immunofluorescent analysis of the TA muscles from Myf6-KO and WT animals with antibodies against Pax7 as a marker of MuSCs and Laminin to mark the periphery of myofibers. Our data indicates that loss of Myf6 has no significant effect on the establishment of MuSC pool during development as the MuSC compartment remains unchanged at postnatal day 7 (P7) and P21 (Fig. 5a-d). However, by the age of three months we observed a significant reduction in the number of Pax7^+^ cells in the Myf6-KO TA muscles compared to their WT counterparts (Fig. 5e-f). The reduction in the stem cell pool continues during late adulthood at 10 months of age (Fig. 5g-h). Likewise, analysis of muscle stem cells associated with the Extensor Digitorum Longus (EDL) muscles stained with anti-Pax7 antibody immediately after isolation (T=0 hour post isolation) showed a similar significant reduction in the number of MuSCs in Myf6-KO myofibers (Fig. i-l). Taken together this data suggests that maintenance of the MuSC pool in adult mice is dependent of Myf6.

**Figure 5.**
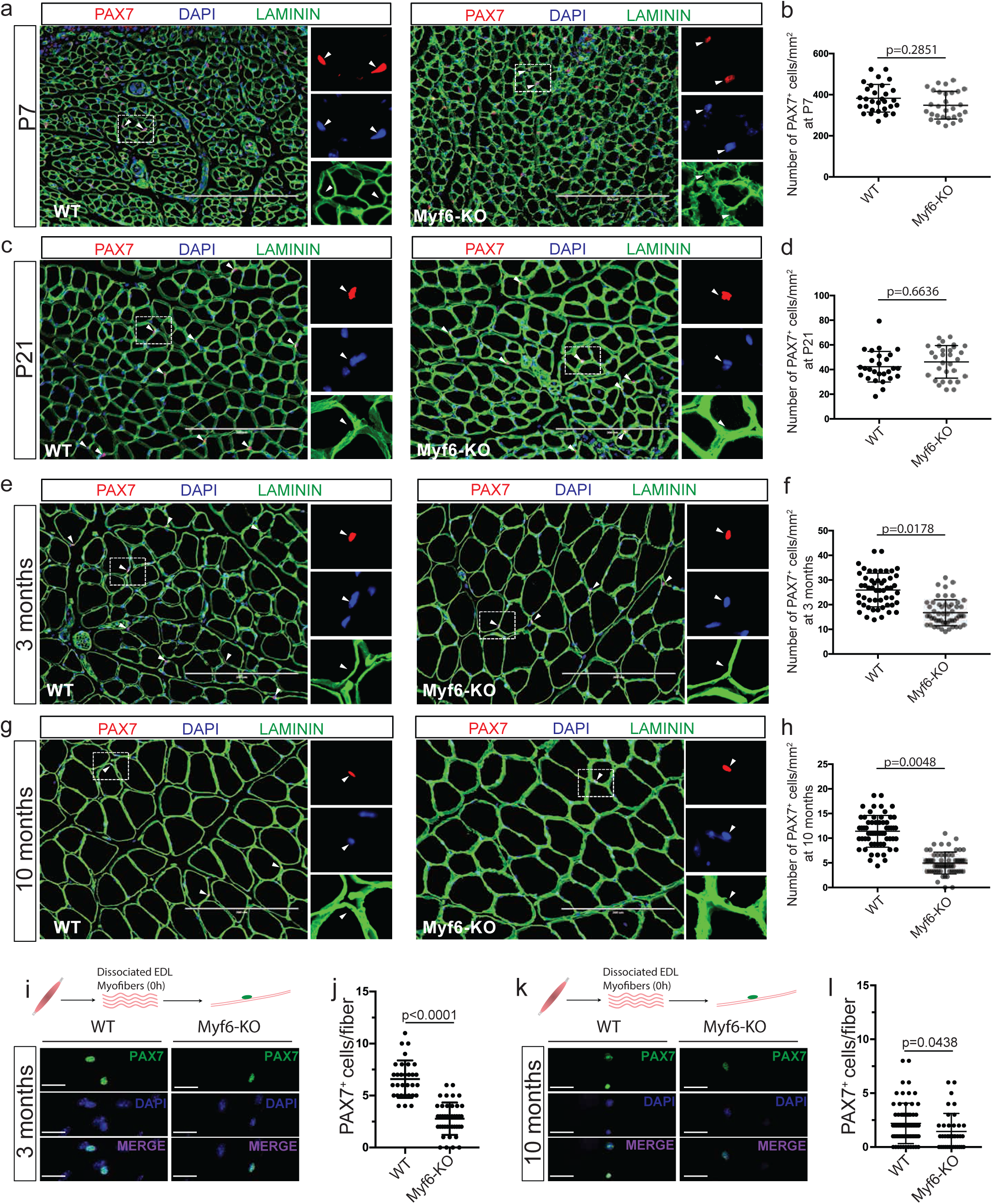
Genetic Deletion of Myf6 Leads to Depletion of the Muscle Stem Cell Pool. **(a)** Immunofluorescent analysis of TA muscle cross-sections from Myf6-KO and WT mice with PAX7 and Laminin (LAMA1) antibodies at postnatal day 7. **(b)** Quantification of the number of PAX7^+^ cells per unit area (mm^2^) in TA muscle cross-sections from Myf6-KO and WT counterparts at postnatal day 7. (n=3 animals per group). **(c)** Immunofluorescent analysis of TA muscle cross-sections from Myf6-KO and WT mice with PAX7 and Laminin (LAMA1) antibodies at postnatal day 21. **(d)** Quantification of the number of PAX7^+^ cells per unit area (mm^2^) in TA muscle cross-sections from Myf6-KO and WT counterparts at postnatal day 21. (n=3 animals per group). **(e)** Immunofluorescent analysis of TA muscle cross-sections from Myf6-KO and WT mice with PAX7 and Laminin (LAMA1) antibodies at 3 months of age. **(f)** Quantification of the number of PAX7^+^ cells per unit area (mm^2^) in TA muscle cross-sections from Myf6-KO and WT counterparts at 3 months of age. (n=5 animals per group). **(g)** Immunofluorescent analysis of TA muscle cross-sections from Myf6-KO and WT mice with PAX7 and Laminin (LAMA1) antibodies at 10 months of age. **(h)** Quantification of the number of PAX7^+^ cells per unit area (mm^2^) in TA muscle cross-sections from Myf6-KO and WT counterparts at 10 months of age. (n=3 animals per group). **(i)** Immunofluorescent analysis of myofiber-associated MuSCs isolated from the EDL muscles of 3 month-old Myf6-KO and WT counterparts. Fibers were stained with PAX7 antibody and counterstained with DAPI at time T=0 hours post fiber isolation. **(j)** Quantification of the number of MuSCs per fiber between 3 month-old Myf6-KO and WT counterparts (n=3 animals per group, p-values are based on two tailed t-test). **(k)** Immunofluorescent analysis of myofiber-associated MuSCs isolated from the EDL muscles of 10 month-old Myf6-KO and WT counterparts. Fibers were stained with PAX7 antibody and counterstained with DAPI at time T=0 hours post fiber isolation. **(l)** Quantification of the number of MuSCs per fiber between 10 month-old Myf6-KO and WT counterparts (n=3 animals per group, p-values are based on two tailed t-test).

### Deletion of Myf6 Leads to Spontaneous Break From Quiescence and Entry of MuSCs into Differentiation

To determine the mechanism underlying the exhaustion of the muscle stem cell pool in Myf6-KO adult mice we first examined the quiescent state and the size of the muscle stem cell pool during three postnatal time points. Immunofluorescence analysis of TA muscle at postnatal day 7 (P7) and P21 indicates that the size of the stem cell pool is equal between Myf6-KO and WT counterparts during early postnatal life (Fig. 5a-d, Fig. S4a-d, Fig. 6a). However, by the third week of postnatal life (P21), muscle stem cells from the Myf6-KO animals spontaneously break quiescence as shown by increased number of Pax7^+^ cells that also express KI67, a marker for proliferating cells (Fig. 6c-f). The spontaneous break of MuSCs from a quiescent state continues postnatally and by three months of age there is a significant reduction in the MuSC pool of Myf6-KO animals compared to their WT counterparts (Fig. 5e-f, Fig. 6e-f). Next, isolated MuSCs from the Myf6-KO and their WT counterpart by FACS using ITGA7^+^/Lin^−^, as described in materials methods, and performed *in vitro* analysis of MuSCs from the Myf6-KO and WT counterparts for differentiation potential, senescence and apoptosis. Our data indicates that loss of Myf6 has no significant effect on the propensity of MuSCs to undergo apoptosis or senescence (Fig. S4e-h), suggesting that neither apoptosis nor cellular senescence are the cause of MuSC exhaustion in Myf6-KO muscle. Next, we performed immunofluorescence analysis of myofiber-associated MuSCs from the EDL myofibers at T0 post fiber isolation. At T0, MuSCs express PAX7 but not MYOD. However, in Myf6-KO EDL myofibers we observed a significant number of MuSCs that express both PAX7 and MYOD, suggesting that a significant portion of MuSCs from the Myf6-KO mice have exited quiescence (Fig. 6k-i). This data is consistent with our earlier analysis showing that by three weeks of age, MuSCs from Myf6-KO mice break quiescence and express KI67 (Fig. 6c-f). Next, we performed RNA-Seq analysis of freshly sorted MuSCs from Myf6-KO and WT mice. To overcome the limitation in the number of MuScs that can be isolated from each animal and, by proxy, the limited amount of total RNA, we used Switching Mechanisms at 5’ end of RNA Transcript (SMART technology, Clonetech). Using adjusted p-value of <0.01, we identified 754 genes that are significantly up or down regulated in Myf6-KO compared to their WT counterparts (Table S4). Analysis of RNA-Seq data shows that the p38 MAPK pathway is significantly upregulated in MuSCs of the Myf6-KO compared to their WT counterparts (Fig. 6g-h). Up regulation of p38 MAPK was further validated by immunofluorescence analysis of EDL myofibers associated MuSCs (Fig. 6n-o). p38 MAPK plays a critical role in MuSCs differentiation. Notably, our data shows that progenitors derived from MuSCs from Myf6-KO mice readily undergo myogenic differentiation with a fusion index approaching 100% as compared to their WT counterpart which exhibit a fusion index of approximately 60%. MuSCs associated with EDL myofibers from the Myf6-KO mice but not their WT counterparts express MYOD and enter myogenic differentiation program. Taken together, this data suggest that loss of Myf6 in skeletal muscle leads to precocious activation of differentiation program by dampening EGFR and STAT3 pathways and concomitant activation of p38 MAPK (Fig. 4, Fig. 6) and expressing MYOD (Fig. 6k-m).

**Figure 6.**
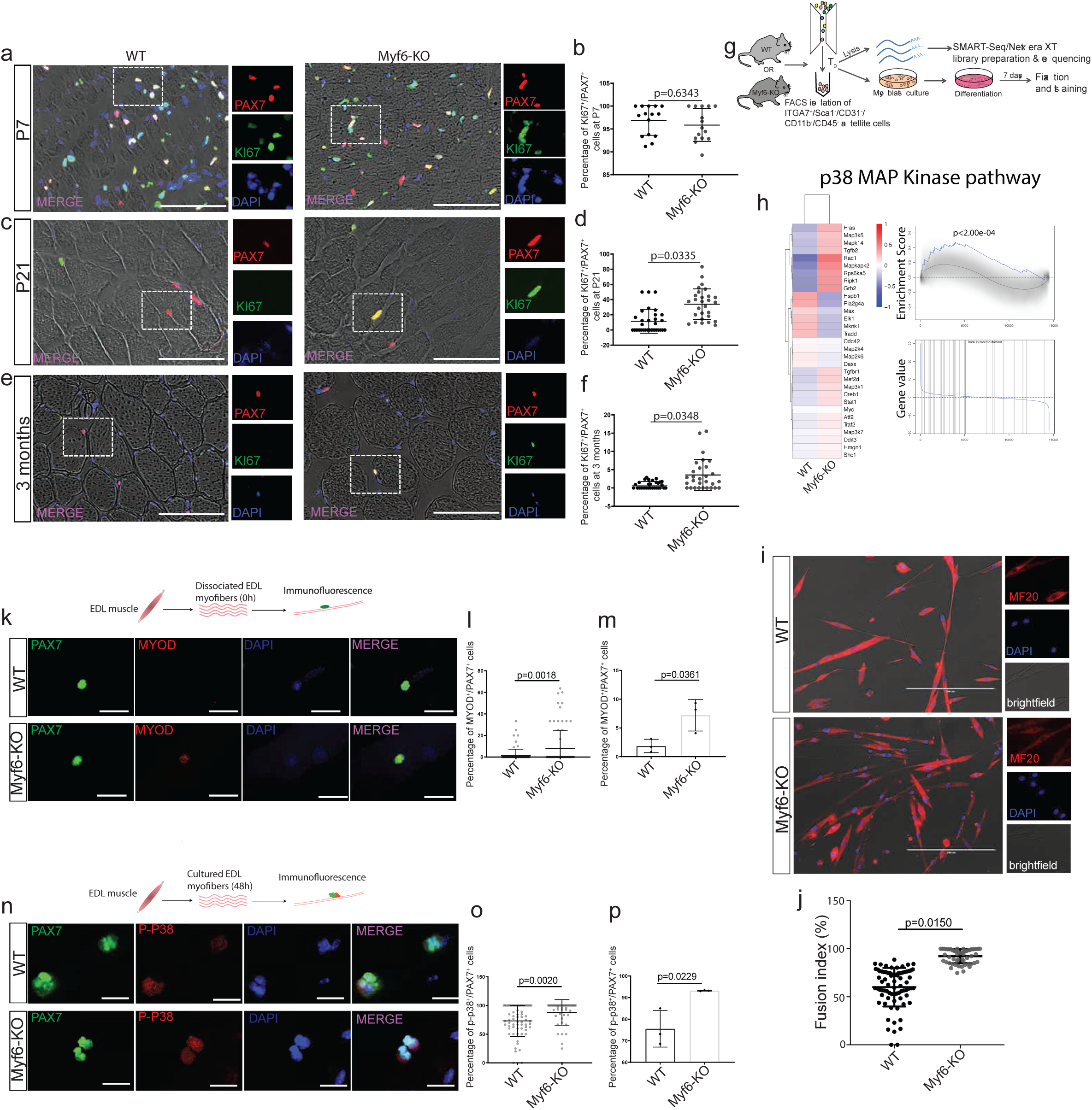
MuSCs from Myf6-KO Mice Exhibit Precocious Exit from Quiescence. **(a)** Immunofluorescent analysis of TA muscle cross-sections from Myf6-KO and WT mice with PAX7 and KI67 antibodies at postnatal day 7. **(b)** Quantification of the percentage of KI67^+^ over PAX7^+^ MuSCs between Myf6-KO and WT mice at postnatal day 7 (n=3 animals per group). **(c)** Immunofluorescent analysis of TA muscle cross-sections from Myf6-KO and WT mice with PAX7 and KI67 antibodies at postnatal day 21. **(d)** Quantification of the percentage of KI67^+^ over PAX7^+^ MuSCs between Myf6-KO and WT mice at postnatal day 21 (n=3 animals per group). **(e)** Immunofluorescent analysis of TA muscle cross-sections from Myf6-KO and WT mice with PAX7 and KI67 antibodies at 3 months of age. **(f)** Quantification of the percentage of KI67^+^ over PAX7^+^ MuSCs between Myf6-KO and WT mice at 3 months of age (n=5 animals per group). **(g)** Schematic drawing of the strategy for FACS isolation of Myf6-KO and WT satellite cells for RNA-Seq library preparation or for culture and differentiation. **(h)** (Left) Heat Map of up and down-regulated genes in the p38 MAPK pathway (WP400) in Myf6-KO and WT satellite cells. (Right) Gene set enrichment analysis of p38 MAPK pathway genes. In each plot, the blue curve represents the gene set enrichment score profile ^37^ across MAPK pathway genes that are ranked by their differential expression. The grey curve represents the null expectation. P-values are based on permutation test (10,000 permutations). **(i)** Immunofluorescent analysis of WT and Myf6-KO primary myotubes cultured in differentiation media for 7 days and stained with MF20 and DAPI. **(j)** Quantification of the fusion index between Myf6-KO and WT 7DM myotubes. Fusion index is calculated as the percentage of myonuclei out of the total number of nuclei per field of view. **(k)** EDL myofibers isolated from WT and Myf6-KO mice fixed at time T=0 and stained with PAX7, MYOD and DAPI. **(l)** Quantification of the average percentage of MYOD^+^/PAX7^+^ satellite cells per mouse in Myf6-KO and WT myofibers (n=3). **(m)** Quantification of the percentage of MYOD^+^/PAX7^+^ satellite cells per fiber in Myf6-KO vs WT myofibers. **(n)** Staining for phospho-p38 MAPK on EDL myofibers cultured for 48 hours from Myf6-KO and WT mice. **(o)** Quantification of the average percentage of phospho-p38^+^/PAX7^+^ satellite cells per mouse in Myf6-KO vs WT myofibers (n=3). **(p)** Quantification of the percentage of phospho-p38^+^/PAX7^+^ satellite cells per fiber in Myf6-KO vs WT myofibers.

### Myf6 is Required for the Establishment of Functional Niche Microenvironment in Skeletal Muscle

Loss of Myf6 led to a reduced MuSC pool in adult mice (Fig. 5a-l) through spontaneous MuSC break from quiescence (Fig. 6a-f). MuSCs from Myf6-KO mice upregulate p38 MAPK pathway, express MyoD (Fig. 6k-p) and enter into myogenic differentiation (Fig. 6g-j). Therefore, we investigated whether defects in the stem cell compartment in Myf6-KO mice are the result of defective niche microenvironments. Since our earlier analysis established the absolute requirement for Myf6 to regulate the expression of key myokines (Fig. 3-4), resulting in impaired activation of EGFR and STAT3 (Fig. 4d-g), we hypothesized that genetic deletion of Myf6 leads to a defective niche microenvironment which is pro differentiation. To test this hypothesis, we isolated MuSCs from a luciferase mouse (B6;FVB-*Ptprc^a^* Tg(CAG-luc,-GFP)L2G85Chco *Thy1^a^*/J) (see Materials and Methods) and transplanted ten thousand cells into either homozygous Myf6 knockout or WT control. Next, we monitored the engraftment efficiency by bioluminescence imaging at day 1, 7, 14 and 21. Notably, in the Myf6 knockout animals there is significantly less retention of donor MuSCs (Fig. 7a-d), suggesting that the main effect of Myf6 on the MuSC compartment is due to a defective niche microenvironment resulting from loss of Myf6. To eliminate the possibility that defects in the MuSC compartment in Myf6-KO animals might be due to a cell intrinsic defect, we performed acute muscle regeneration assay using CTX injury. Importantly, following CTX-induced injury, the MuSC pool in Myf6-KO mice is reconstituted back to the level seen in their WT counterparts (Fig. 7e-g), suggesting that exposure of muscle stem cells to exogenous source of cytokines from infiltrating immune cells during muscle injury can rescue defects in the MuSC compartment. Similarly, continuous exposure of culturing EDL myofibers with chick embryo extract expands the number of MuSCs associated with EDL myofibers (Fig. 7h-i). Furthermore, exposure of myofiber-associated MuSCs to ligands such as EGF and LIF significantly increases the number of high PAX7 expressing MuSCs (Fig. 7j-k). Lastly, our data shows that cultured MuSCs from Myf6-KO mice have similar growth characteristics as those of their WT counterparts (Fig. 7i-m). Taken together, our data indicates that the defect in the stem cell compartment of the Myf6-KO animals is not due to cell-intrinsic defects but is primarily due to an impairment in the niche microenvironment, in which depletion of myokine expression from the myofiber leads to activation of a signaling cascade that breaks MuSC quiescence, activates p38 MAP and induces precocious differentiation. This perturbation in the niche environment leads to a gradual reduction in the number of MuSCs in adult mice.

**Figure 7.**
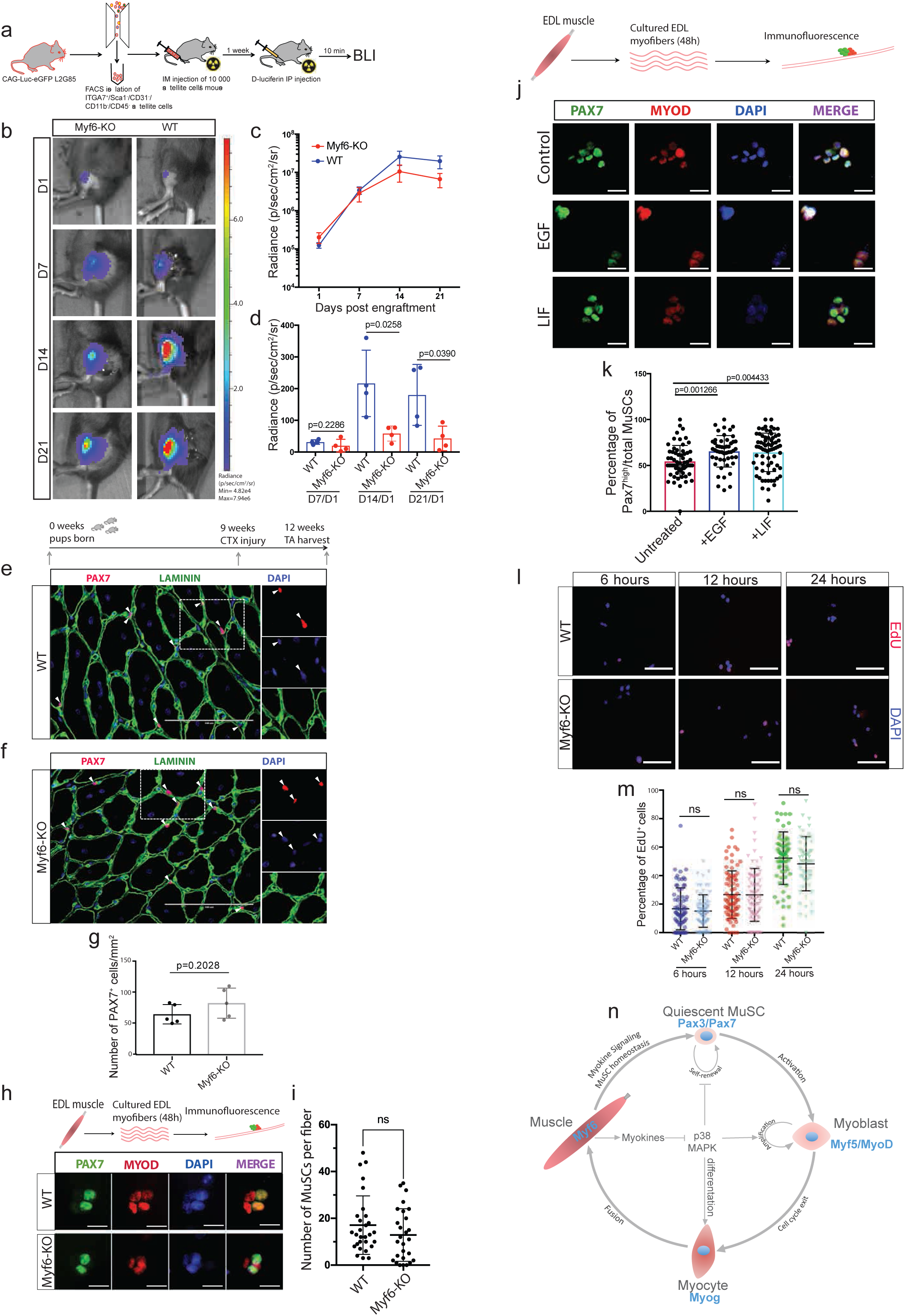
Exposure of MuSCs from Myf6-KO Mice to Exogenous Sources of Cytokines Rescues their Self-Renewal Defects. **(a)** Schematic drawing of MuSC transplantation and *in vivo* bioluminescence experiments. Briefly, 10 000 MuSCs isolated by FACS from CAG-Luc-eGFP mice were injected into the irradiated hindlimb of immunocompromised WT or Myf6-KO mice. Bioluminescent imaging was performed at day 1, 7, 14 and 21 post donor MuSC injections. **(b)** Representative images of bioluminescent signal from hindlimbs of Myf6-KO and WT counterparts at day 1, 7, 14, and 21 post transplantation. **(c)** Time course of average bioluminescent signal (total flux in radiance) in Myf6-KO and WT mice at 1, 7, 14 and 21 days post MuSC transplantation showing engraftment dynamics of donor MuSCs (mean across 4-5 animals per group ± SEM at each time point). **(d)** Bioluminescent signal (radiance) in Myf6-KO and WT mice at each time point normalized to the bioluminescent signal at day 1 post MuSC transplantation. (n=4-5 animals per group ± SD) **(e-f)** Immunostaining of TA cross-sections from WT **(e)** and Myf6-KO **(f)** mice stained with laminin (LAMA1) and PAX7 antibodies. Acute muscle injury was performed by injection of Cardiotoxin (CTX) into the TA muscle of Myf6-KO and WT mice (n=5 animals per group). **(g)** Quantification of the number of PAX7^+^ cells per unit area of TA muscle cross sections between Myf6-KO and WT mice (n= 5 animals per group, p-value is based on two tailed t-test). **(h)** Immunofluorescent analysis of Myf6-KO and WT EDL myofibers cultured in growth media for 48 hours and stained with PAX7 and MYOD antibodies. **(i)** Quantification of the number of PAX7^+^ MuSCs per fiber in Myf6-KO and WT mice after 48 hours in culture. **(j)** Immunofluorescence staining of WT EDL myofibers cultured in growth media for 48 hours and treated with LIF (1000U/ml) or EGF (80ng/ml) every 8 hours. Fibers were stained with antibodies against PAX7 and MyoD. **(k)** Quantification of the number of PAX7^+^ and MYOD^+^ cells on the EDL myofiber treated with recombinant LIF, EGF or vehicle control. **(l)** EdU staining of cultured primary myoblasts isolated by FACS (see materials and methods) from Myf6-KO and WT mice. Staining was analyzed after 6, 12 and 24 hours of EdU incorporation. **(m)** Quantification of the percentage of EdU^+^ myoblasts after 6, 12, and 24 hours of EdU incorporation. **(n)** Schematic drawing of the mechanism of Myf6 action in maintaining MuSC homeostasis. Briefly, myofibers expressing Myf6 secrete myokines that block the activation of the p38 MAPK pathway, which is necessary for maintenance of the MuSC pool.

## Discussion

Skeletal muscle has an inherent endocrine function in addition to its primary functions of producing force and regulating energy metabolism ^21,33,34^. However, mechanistic insights into how myokines are regulated and whether or not they function as part of the MuSC niche milieu remains largely unexplored. Here, we report that Myf6 is critical to maintain the expression of many myokines with integral functions in the control of various signaling pathways in skeletal muscle. Our data identifies a novel function for Myf6 as a myogenic regulator of niche factors, namely myokines (Fig. 1-4).

Consistent with previous work, the differentiation function of Myf6 in myogenesis is dispensable, likely because of the large overlap in the regulatory function of myogenic factors (Fig. S5) ^35^. We found that mice with a homozygous *Myf6* deletion showed no visible abnormalities in skeletal muscle fiber morphology (Fig. S3). The differentiation function of Myf6 is likely compensated redundantly by other myogenic factors, i.e. MyoD and/or Myogenin during transient differentiation of muscle stem cells (Fig. S5). Indeed, by comparative ChIP-Seq analysis we found that Myf6 and MyoD share 50% overlap in their targets during differentiation of muscle cells (Fig. S5).

The unanticipated function of Myf6 in regulating MuSC pool via myokine expression is primarily due to its temporal position in the hierarchy of the muscle differentiation program. While Myf5, MyoD and Myogenin are transiently expressed during MuSCs differentiation, in mature skeletal muscle fibers, Myf6 is the only factor with significant expression (Fig. 1a). Consistent with this notion, comparative analysis of ChIP-Seq and gene expression data in differentiating myotubes shows that MyoD is fully capable of binding to a very similar set of genes, including many genes that are targets of Myf6 (Fig. 2a, Fig. S5).

Our ChIP-Seq (Fig. 1), gene expression analysis(Fig. 2) and ELISA experiments (Fig. 3) point to a role for Myf6 in the production of multiple regulatory myokines such as EGF, VEGFA and LIF. Our striking observation that loss of Myf6 leads to significant reduction of myokines not only in the secretome of differentiating myotubes (Fig. 3f-g) but also in serum (Fig. 3h-i) suggests that the regulatory function of Myf6 may extend beyond skeletal muscle.

The impairment in the ability of MuSCs to activate EGFR and STAT3 followed by subsequent rescue (Fig. 4d-g), strongly suggests that Myf6 regulates MuSC activity by myokine signaling. A critical factor that is required for the maintenance of the MuSC pool throughout adult life is the ability of stem cells to remain quiescent under homeostatic conditions. In Myf6-KO animals, muscle stem cells break quiescence and spontaneously enter into differentiation program by upregulating p38 MAPK pathway and express MYOD (Fig. 6). Such spontaneous exit from quiescence and entry into myogenic differentiation results from abrogation of myokine-mediated communication between myofibers and their associated stem cells in the Myf6-KO animals. The inability of donor stem cells to efficiently engraft into the skeletal muscles of Myf6-KO further supports our conclusion that loss of Myf6 leads to defective communication between MuSCs and their niche microenvironment. Our *in vivo* and *in vitro* analysis uncover a novel Myf6-mediated myokine signaling pathway that establishes a direct communication between myofibers and their associated stem cells. Therefore, we propose that Myf6 is a myogenic niche regulator which acts upstream of EGFR and STAT3 pathways to maintain muscle stem cell quiescence and maintain muscle stem cell pool.

## Acknowledgements

We thank Christian Young at the Lady Davis Institute for Medical Research – Jewish General Hospital – core facility for help with the Fluorescence Activated Cell Sorting (FACS). We thank Dr. Colin Crist at the Lady Davis Institute and McGill University Department of Human Genetics for sharing reagents. This work was supported by a grant from the Canadian Institute of Health Research (CIHR) project grant PJT-156087 to VDS, an NSERC discovery grant to VDS, a research grant from the Richard and Edith Strauss Foundation to VDS and by the Canada Research Chairs Program (CRC) to VDS.

## Authors Contribution

Conceived the idea, VDS; performed experiments, VDS, FL, DB, KS, NK, DN, CL; Analyzed data, VDS, FL, DN, TJP, HSN, AHC; wrote the manuscript; VDS, FL.

## Materials and Methods

### KEY RESOURCES TABLE

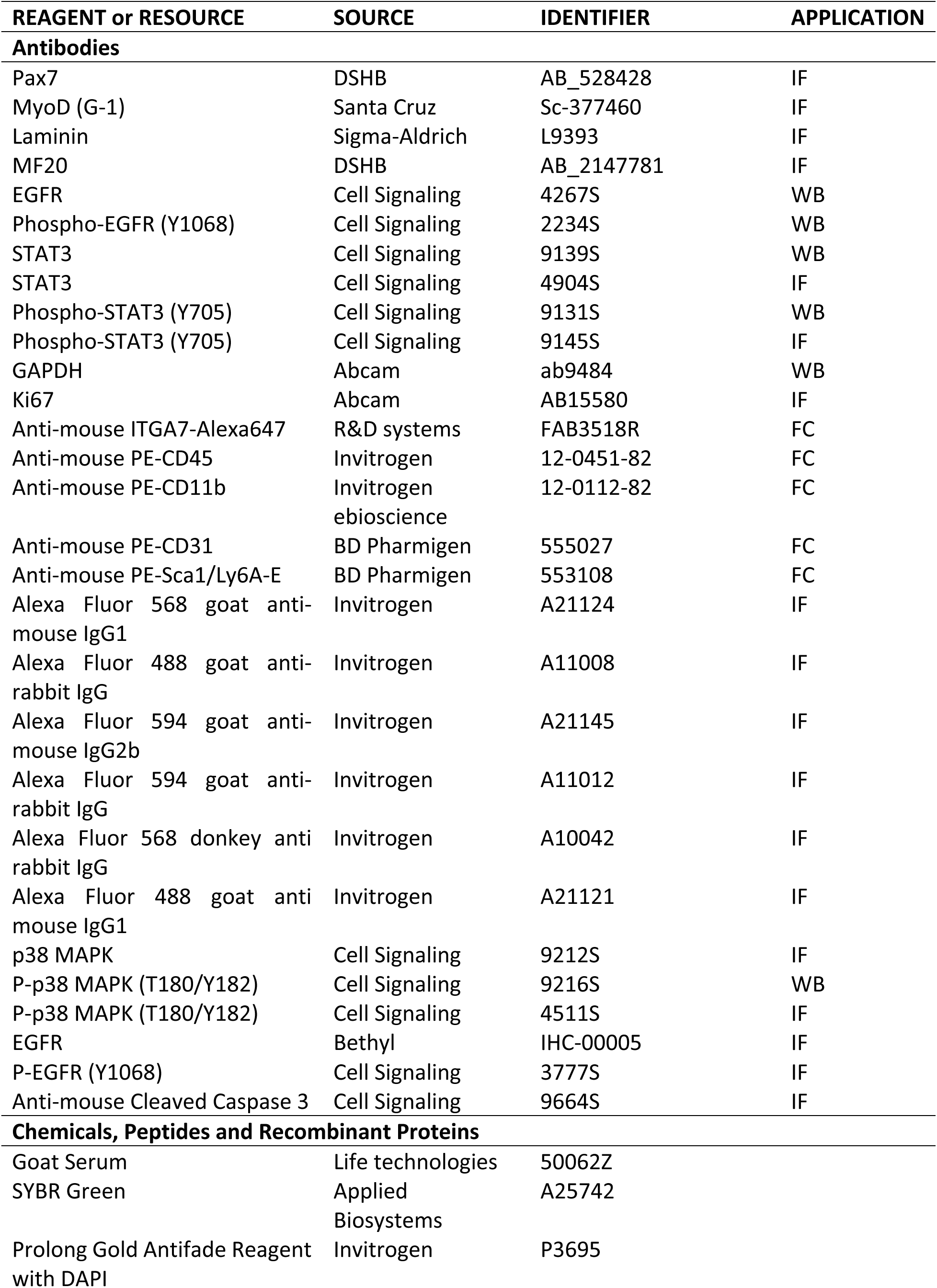
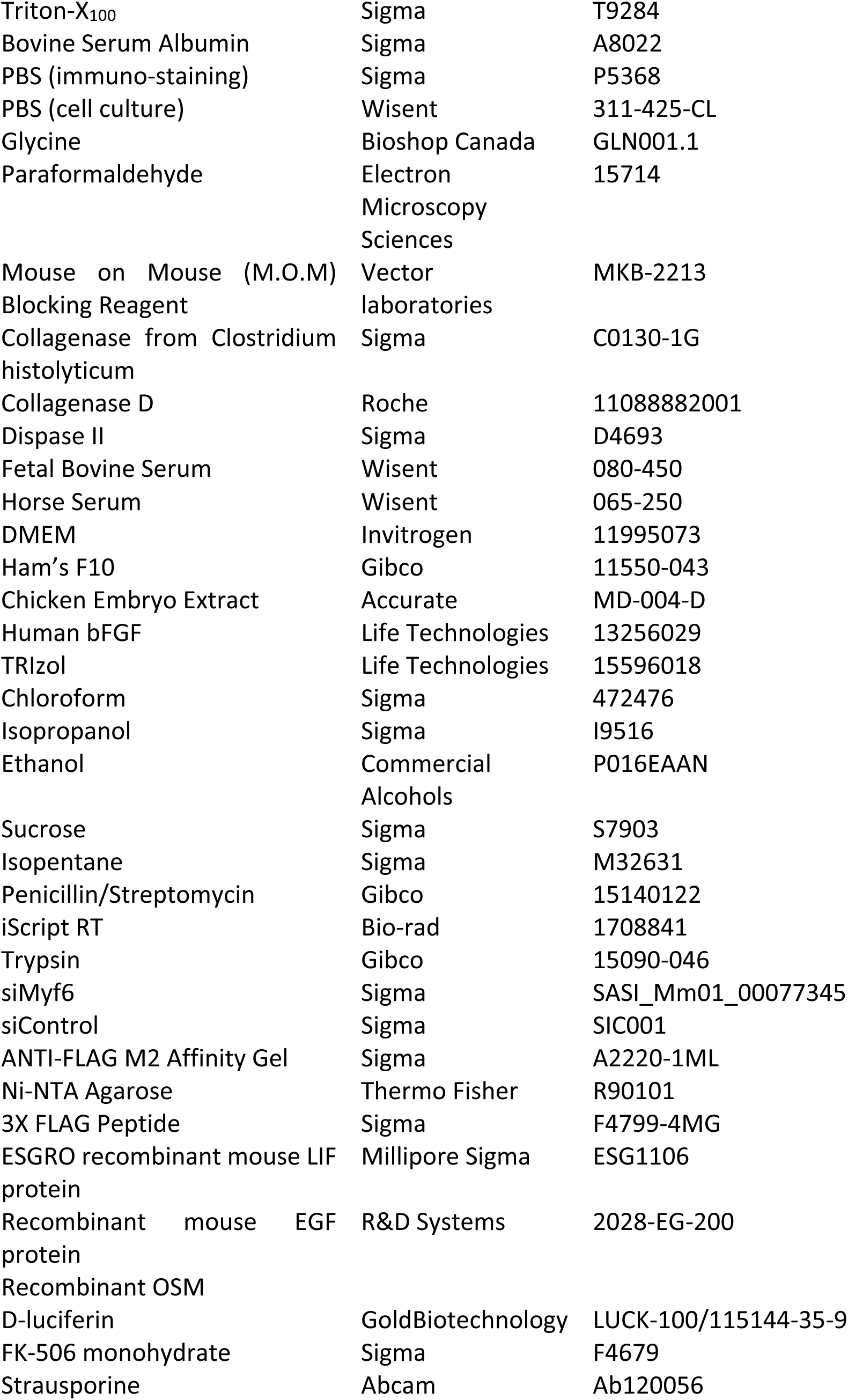
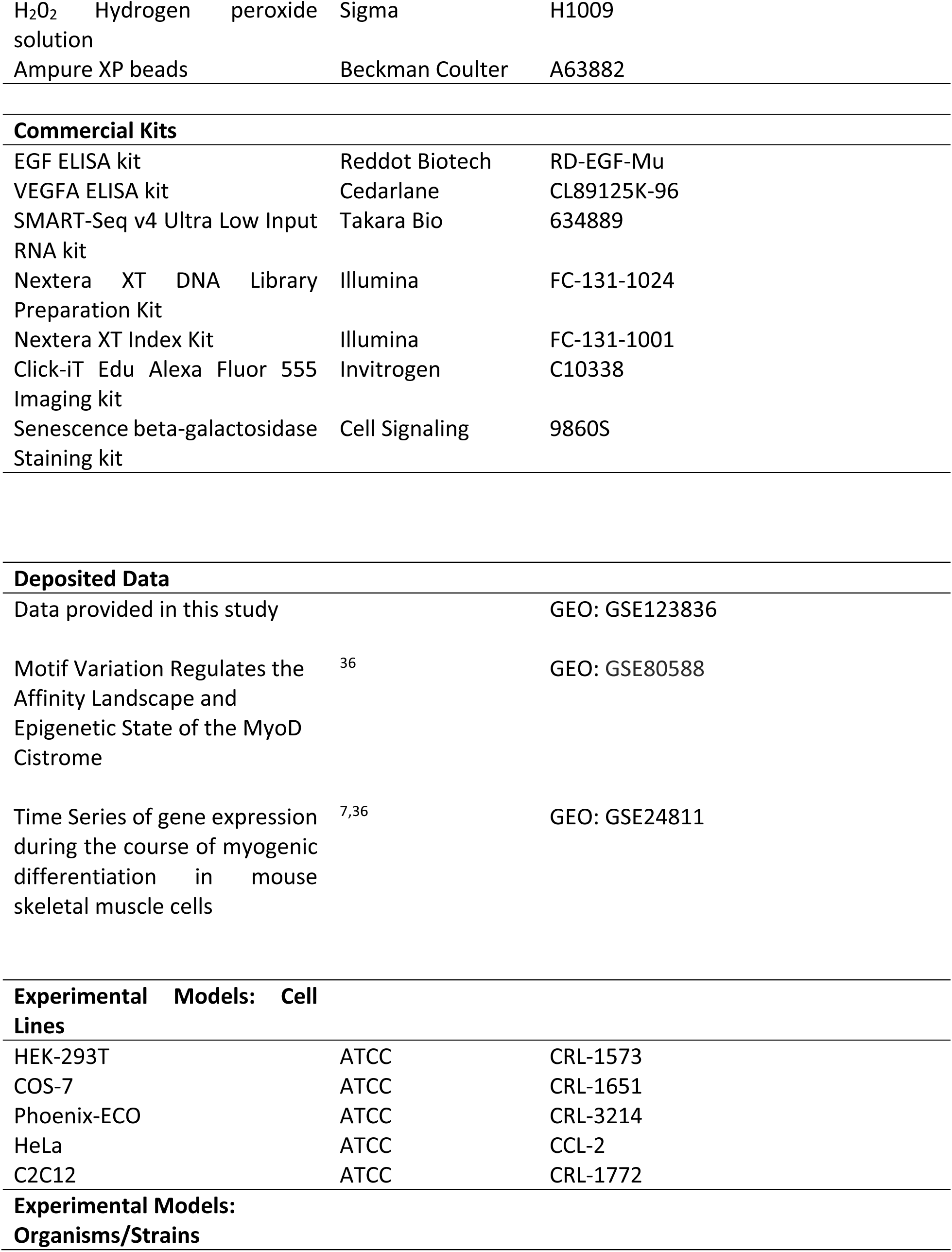
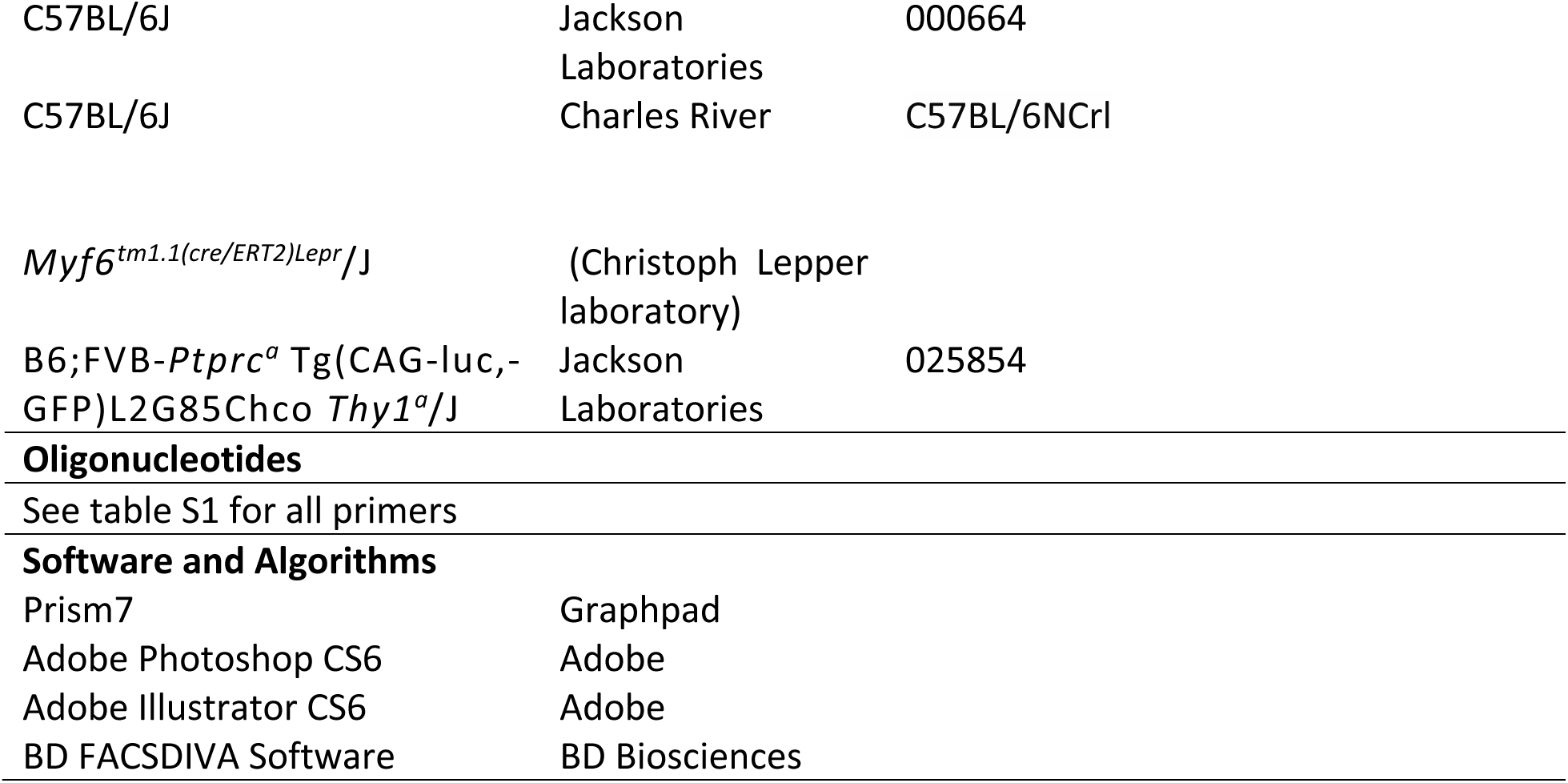

### CONTACT FOR REAGENT AND RESOURCE SHARING

Further information and requests for resources and reagents should be directed to and will be fulfilled by the Lead Contact, Vahab Soleimani

### EXPERIMENTAL MODEL AND SUBJECT DETAILS

#### Animal Procedures

All animal procedures were approved by the McGill University Animal Care Committee (UACC).

#### Cell Isolation and Culture

Muscle stem cells were isolated from the hindlimb muscles of 8-week-old mice by Fluorescence Activated Cell Sorting, as described in the Method Details. Sorted cells were cultured on collagen-coated plates in growth media (Ham’s F10 supplemented with 20% FBS, 1% penicillin streptomycin and 2.5 ng/ml bFGF) and passaged after reaching 60% confluency.

### METHOD DETAILS

#### Generation of Myf6-CTAP Retrovirus

Full Open reading Frame (FL-ORF) of the mouse Myf6/MRF4 was sub cloned into pBRIT retroviral backbone ^6^. The construct was verified by sequencing and its functionality was confirmed using multiple functional assays (Fig. S1). Retroviral particles containing Myf6-CTAP were generated by transfection of Phoenix helper-free retrovirus producer lines, a kind gift from the Nolan lab (https://web.stanford.edu/group/nolan/_OldWebsite/retroviral_systems/phx.html) with Polyethylenimine (PEI, Polysciences Cat. 23966-2). Viral supernatant was collected after 48 hours post transfection.

#### Generation of Primary Myoblasts Over-Expressing Myf6-CTAP

Muscle stem cells were isolated from 8-week old WT C57BL/6 mice by FACS and were propagated and maintained in growth media (HAM’s F10 supplemented with 20% FBS, 1% penicillin/streptomycin and 2.5 ng/ml basic Fibroblast Growth Factor, bFGF) for up to three passages. The resulting myoblasts were then infected with a retroviral cocktail containing Myf6-CTAP virus and polybrene (Millipore) for 8 hours. Cells were subsequently washed with 1X PBS and fed with normal growth media. The selection of transduced cells was carried out by puromycin selection (2.5 μg/ml) in primary myoblast growth media for up to one week. Subsequently, stably transduced primary cells were maintained in low puromycin selection media (HAM’s F10 supplemented with 20% FBS, 2.5 ng/ml bFGF, 1% penicillin/streptomycin, and 1.25 μg/ml puromycin). Differentiated multinucleated myotubes were obtained by placing >90% confluent myoblasts in differentiation media (DMEM supplemented with 5% horse serum) for five days.

#### Tandem Affinity Chromatin Immunoprecipitation Sequencing (ChTAP-Seq)

ChTAP-Seq was carried out as described previously ^26^. Briefly, multi nucleated myotubes over expressing Myf6-CTAP or empty vector- (EV-CTAP) were harvested and 20 mg of chromatin was used as input for ChIP. ChIP was performed in tandem using anti FLAG antibody conjugated to agarose beads (Sigma). Chromatin was then eluted from the beads by competition with FLAG peptide (Sigma) and cleavage by TEV protease (New England Biolabs). The eluted chromatin was then used for a second purification step using Ni-NTA Agarose with nickel-charged affinity resin (Thermo Fisher), as described previously ^26^. Finally, 10 ng of ChIP DNA was used to construct ChIP-Seq libraries using TruSeq ChIP Library Preparation Kit (Illumina). Reads from the Myf6 ChTAP- and the EV-ChTAP-Seq were mapped to the mouse mm10/GRCm38 mouse genome assembly. Peak calling was carried out using MACS2.0 ^27^ with p-value threshold of 10^−5^ using default parameters.

#### Luciferase Assays

Luciferase assays were carried out as described previously ^36^. Briefly, Cos7 cells were co transected with retroviral vectors harboring MRFs (MyoD or Myf6) with and without mouse E47 expression vector together with the luciferase vector (pGL4.23 [luc2/minP]) containing a multimerized E-box motif with three spaced out CAGCTG motifs, as described previously ^36^. Dual luciferase assay was performed using Promega Dual luciferase assay kit. Relative luciferase expressions were normalized to renilla and plotted relative to the promoter readout.

#### Immunofluorescent Analysis of the Tibialis Anterior (TA) Muscles

TA muscle sections were fixed in 4% paraformaldehyde (PFA) in PBS for 10 minutes on ice and washed 3 times for 5 minutes each in PBS. Permeabilization was done in 0.3% Triton-X_100_ and 0.1M glycine in PBS for 20 minutes on a shaker at room temperature, followed by washing once in PBT (0.05% Triton-X_100_ in PBS, v/v).

Muscle cross sections were first blocked in Mouse blocking reagent (M.O.M reagents Vector Laboratories) for one hour and subsequently in 10% goat serum (Thermo Fisher) supplemented with 3% Bovine Serum Albumin (BSA) in a humid chamber at room temperature for one hour. Subsequently, muscle sections were incubated with the primary antibodies (Pax7, concentrated DSHB 1:100 dilution; Laminin (LAMA1), Sigma 1:750 dilution) in goat serum with BSA blocking solution overnight in a humid chamber at 4 **°**C. The next day, slides were washed twice for 10 minutes each time in PBT (0.05% Triton-X_100_ in PBS, v/v), and once for 10 minutes in PBS. Samples were then incubated with the secondary antibodies (goat anti-mouse IgG1, Alexa 568 1:1000 dilution; goat anti-rabbit IgG Alexa 488 1:500 dilution) diluted in goat serum in BSA blocking solution for one hour at room temperature in a humid chamber protected from light. Sections were washed three times for 10 minutes each time in PBT followed by mounting with ProLong Gold AntiFade plus DAPI solution (Invitrogen P36935) and coverslips were sealed using clear nail polish.

#### Skeletal Muscle Injury and Regeneration

Injury-induced regeneration experiments were performed via intra-muscular injection of 50 μl cardiotoxin (CTX, 5 μM, Sigma) into the TA muscle of 3-4-months old *Myf6-KO* mutant and WT control animals. Prior to injury, mice were anaesthetized by intra-peritoneal injection of Avertin (2,2,2-tribromoethanol from TCI America at 15 μl per gram body weight of 20 mg per ml solution), or by the administration of Isoflurane. The right injured and the left uninjured TA muscles were harvested via dissection 21 days post CTX injury. Muscles were immediately mounted in tragacanth (Alfar Aesar) on cork and snap-frozen in liquid nitrogen-chilled isopentane (Sigma).

#### Isolation and Immunofluorescent Analysis of EDL Myofibers

Briefly, the skin from the hindlimb of the mouse was removed and the TA muscle was dissected to gain access to the Extensor Digitorum Longus (EDL). Excess tissue was removed around the knee to reveal the proximal tendon of the EDL muscle. The EDL muscle was dissected by cutting the distal tendon, gently peeling the muscle off the leg to reach the proximal tendon and severed that as well. The EDL was placed in a 1.5 ml tube with 1000 U/ml collagenase in Dulbecco’s Modified Eagle’s Medium (DMEM) for one hour at 37 **°**C, 5% CO_2_. The EDL muscle was then transferred to a 6-well plate containing unsupplemented DMEM that had previously been coated with 10% horse serum (HS) plus DMEM for 30 minutes. The fibers were released from the EDL muscle by using a large bore glass pipette with blunted edges at the nozzle, coated with HS, to gently pipette the muscle up and down.

Isolated fibers were transferred to a single well of a 12-well plate that was coated with 10% HS plus DMEM for 30 minutes. Excess media was removed and fibers were fixed with 4% PFA in PBS for 10 minutes at room temperature. After PFA removal, fibers were washed 3X with 0.1% Triton X_100_ in PBS. Fibers were then permeabilized with 0.1% Triton X_100_, 0.1M glycine in PBS for 15 minutes at room temperature (RT). After removal of permeabilization buffer, fibers were washed 3X with 0.1% Triton X_100_ in PBS and blocked with a blocking solution composed of 5% HS, 2% BSA, 1% Triton X_100_ in PBS for at least 1 hour at RT. Subsequently, blocking solution was removed and fibers were stained with appropriate primary antibodies diluted in the blocking solution overnight at 4 °C. The next day, the primary antibody solution was removed and fibers were washed 3X with 0.1% Triton X_100_ in PBS. Subsequently, the secondary antibodies were diluted in the blocking solution and added to fibers at RT for at least one hour. Lastly, antibody solution was removed and fibers were washed 3X with 0.1% Triton X_100_ in PBS and placed on a glass slide with mounting solution containing DAPI for microscopic analysis.

Immunofluorescence of cultured fibers was essentially the same as above with the exception that the fibers were placed in growth media [DMEM/sodium pyruvate (110 mg/L), supplemented with 10% FBS, 1% penicillin/streptomycin, 1% chick embryo extract, 2.5 ng/ml bFGF] and incubated at 37 **°**C and 5% CO_2_ for 48 hours prior to fixation.

#### Immunofluorescence Imaging

Imaging was performed using an EVOS FL Imaging System (Thermo Fischer, AMF4300) equipped with GFP, Texas Red and DAPI filters, and 4x, 10x, 20x, 40x objectives.

#### Quantitative Real-Time PCR

qPCR analysis was performed using SYBR Green qPCR Master Mix containing buffer, dNTPs and Thermo stable hot-start DNA polymerase (Thermo Fisher). Gene-specific primers were designed using GenScript software. (https://www.genscript.com/tools/real-time-pcr-tagman-primer-design-tool) crossing exon junctions for specified genes. The reactions were run on a QuantStudio 7 Flex Real-Time PCR System (Applied Biosystems – Thermo Fisher). Relative expression values were calculated based on the ΔΔCT method using RPS2 or GAPDH as housekeeping genes.

#### Isolation of Muscle Stem Cells by Fluorescence Activated Cell Sorting

Hindlimb muscles from Myf6-KO and WT mice were minced and digested in FACS digestion media, composed of HAM’s F10 with 2.4 U/ml Collagenase D, 2.4 U/ml Dispase II and 0.5 mM CaCl_2_, for 60 minutes at 37**°**C in a 5% CO_2_ Tissue Culture (TC) incubator with intermittent trituration. The digested muscle slurry was supplemented with 9ml of FBS, filtered through a 40μm cell strainer and cells were pelleted by centrifugation at 500g, Relative Centrifugation Force (RCF). The cell pellet was resuspended in 500μl of 2% FBS: PBS (v/v) and incubated with antibodies and live cell stain (Alexa647-conjugated to anti-mouse ITGA7, PE-CD31, PE-CD45, PE-CD11b, PE-SCA1 and Hoechst 33342) for 15 minutes at room temperature. Cells were washed in 2% FBS:PBS (v/v) solution and sorted for the ITGA7^+^/Hoechst^+^/CD31^−^/CD45^−^/CD11b^−^/SCA1^−^ population as described previously ^38^ on a FACSAria Fusion system (Beckman-Dickinson).

Isolated cells were sorted directly into lysis buffer for SMART RNA-Seq library preparation or were cultured in growth media (Ham’s F10 supplemented with 20% FBS, 1% penicillin streptomycin and 2.5 ng/ml bFGF) and passaged after reaching 60% confluency.

#### SMART RNA-Seq Library Preparation

3000 quiescent satellite cells were lysed after FACS by sorting directly into a 0.2ml tube containing 1 μl SMART-Seq Reaction Buffer (95% SMART-Seq 10x lysis buffer containing 5% SMART-Seq RNAse Inhibitor, Takara Bioscience, Cat. 634890) in 8μl ddH_2_0. RNA was reverse transcribed using SMART-Seq Reverse Transcriptase followed by 11 cycles of PCR amplification (SMART-Seq v4 kit, Takara Bioscience, Cat. 634890). Purification and size selection of DNA was carried out using Ampure XP beads (Beckman Coulter, Cat. A63880). Total cDNA was quantified using Quant-iT PicoGreen dsDNA Assay Kit (Thermo Fisher). Sequencing libraries were created with the Nextera XT kit (Illumina Cat. FC-131-1024 and FC-131-2001), using 150 pg of total cDNA as input for tagmentation. Illumina sequencing adapters were added by PCR using 12 cycles of amplification (Illumina Cat. FC-131-1024 and FC-131-2001). Final sequencing libraries were purified and size selected using Ampure XP beads (Beckman Coulter, Cat. A63880).

#### Satellite Cell Transplantation

Hindlimbs of WT and Myf6-KO mice were irradiated with 18Gy using a MultiRad 225 irradiator and were administered a subcutaneous injection of immunosuppressant FK506 (2.5mg/kg) the day before satellite cell transplantation. Luciferase-expressing satellite cells were FACS sorted from B6;FVB-*Ptprc^a^* Tg(CAG-luc,-GFP)L2G85Chco *Thy1^a^*/J mice into 2%FBS:PBS (v/v) and 10 000 cells were injected into the hindlimb of WT and Myf6-KO mice using a Hamilton syringe. FK506 injections (2.5mg/kg) were maintained every 48hours throughout throughout the bioluminescence imaging time-course.

#### *In vivo* Bioluminescence Imaging

Mice were anesthetized and given two 100μl contralateral intraperitoneal injections of D-luciferin (Gold Biotechnology) diluted to 15mg/ml in PBS. Bioluminescence imaging was performed 11 minutes after luciferin injections using an IVIS Spectrum machine (Perkin Elmer). Bioluminescence signal was measured using region of interest (ROI) placed over each hindlimb. Imaging was performed 1, 7, 14 and 21 days after donor cell transplantation.

#### Whole muscle RNA-Seq Preparation

Total RNA was isolated from the hindlimb skeletal muscles of 4-months old Myf6-KO and WT mice using TRIzol reagent (Thermo Fisher). RNA quality and quantity was assessed by the Agilent Bio analyzer and 1.0 μg of RNA was used to construct RNA-Seq libraries. cDNA libraries were constructed from total RNA with the TruSeq RNA v2 kit (Illumina). The size distribution of the libraries was analyzed with the Fragment Analyzer High Sensitivity NGS assay (AATI). DNA concentration was measured with the Qubit High Sensitivity DNA assay (Thermo Fisher).

#### RNA-Seq Analysis

Sequencing was performed on Illumina NextSeq 500 High Output Flow Cell. Sequenced reads were mapped to mouse mm10 genome assembly using HISAT2 ^39^. We used FeatureCounts to quantify gene expression using GENCODE gene definitions ^40^. We then normalized expression by adding one count to every gene (to avoid zeros) and converted gene expression into reads per million (RPM). Differentially expressed genes were identified using the R package DESeq2 ^41^. P-values were corrected by independent hypothesis weighting ^42^.

#### GSEA Pathway Analysis

Genes for p38 MAPK (WP400) pathways were used to perform gene set enrichment analysis ^43-45^ using a weighted Kolmogorov-Smirnov test. In each test, genes in the pathway were weighted by its Wald-test statistic of differential expression. The null distribution of enrichment scores was calculated by permuting the gene labels 10 000 times, the enrichment significance (p-value) was calculated by the comparison of the enrichment score of the original labels with the null distribution ^37^

### QUANTIFICATION AND STATISTICAL ANALYSIS

Statistical analysis was performed using Prism7 (Graphpad). An unpaired two tailed t-test was used for comparison between experimental and control groups. Error bars are representations of the standard deviation unless otherwise stated. Asterisks (*, **, ***, ****) correspond to p-values of <0.05, <0.01, <0.001 and <0.0001, respectively.

### DATA AND SOFTWARE AVAILABILITY

The primary ChIP-Seq and RNA-Seq data reported in this paper is deposited in NCBI and available through Geo Accession number GSE123836.

## Supplemental Information

**Table S.1.**
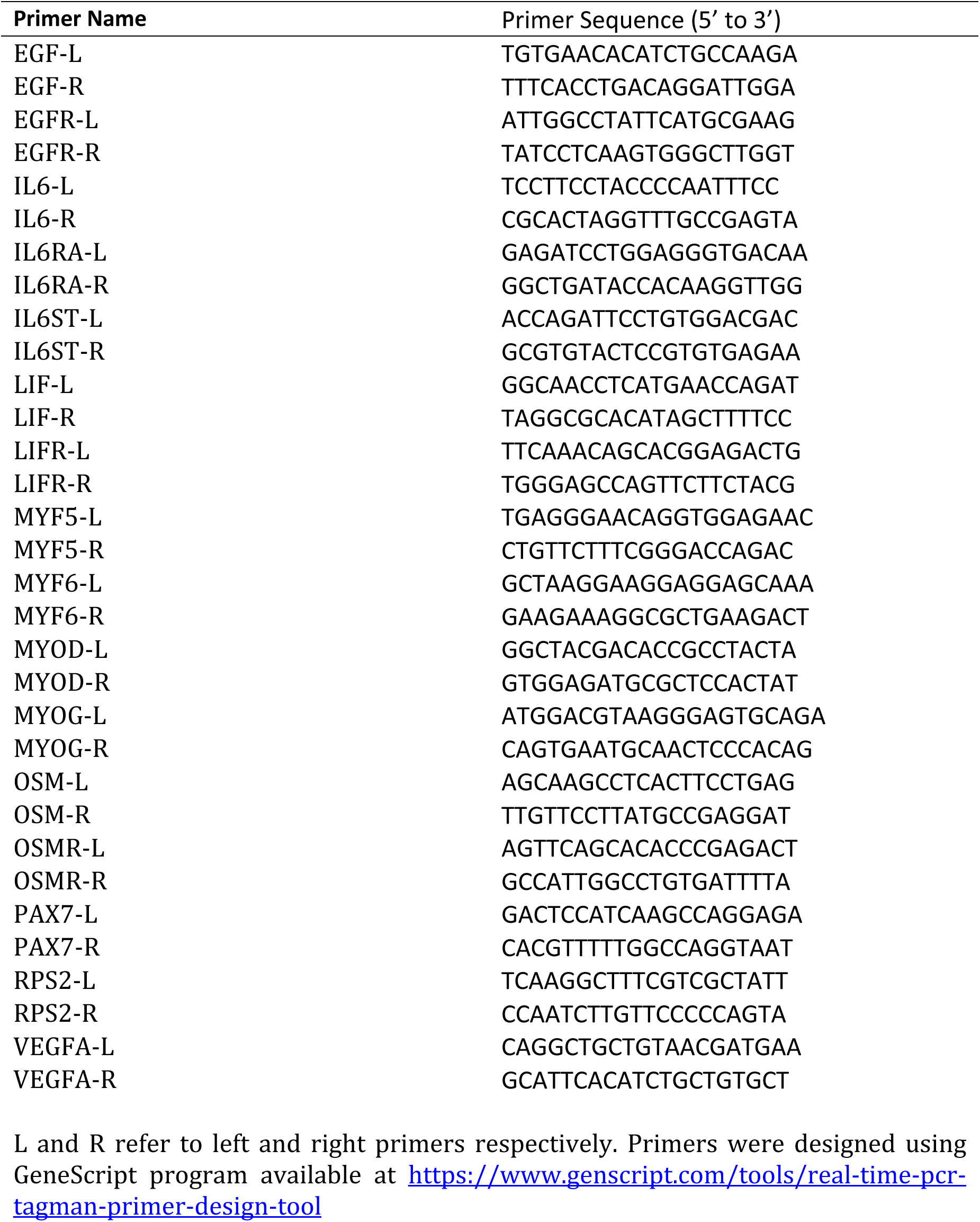
Primer Names and Their Sequences, Used in This Study.

**Table S.2.**
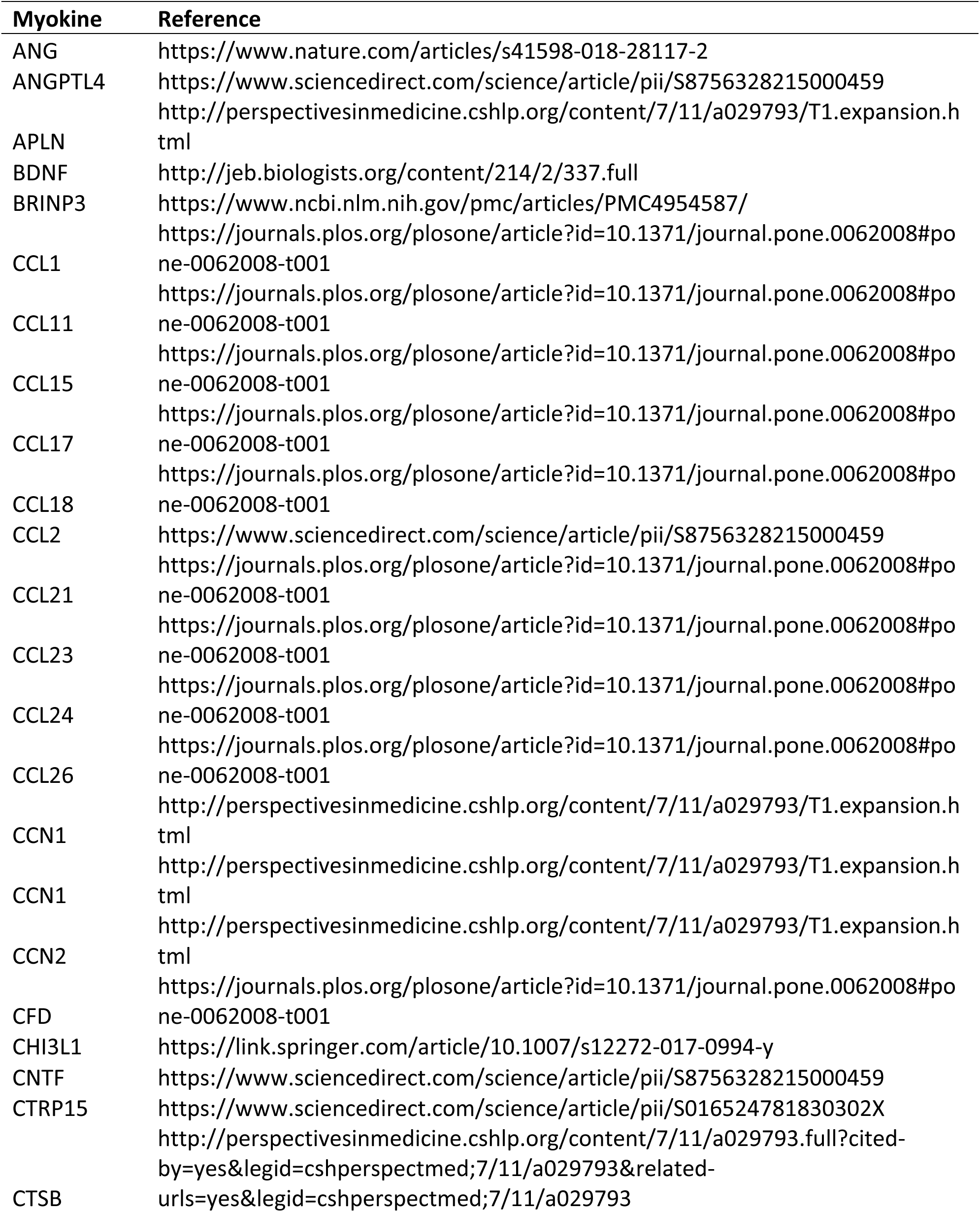
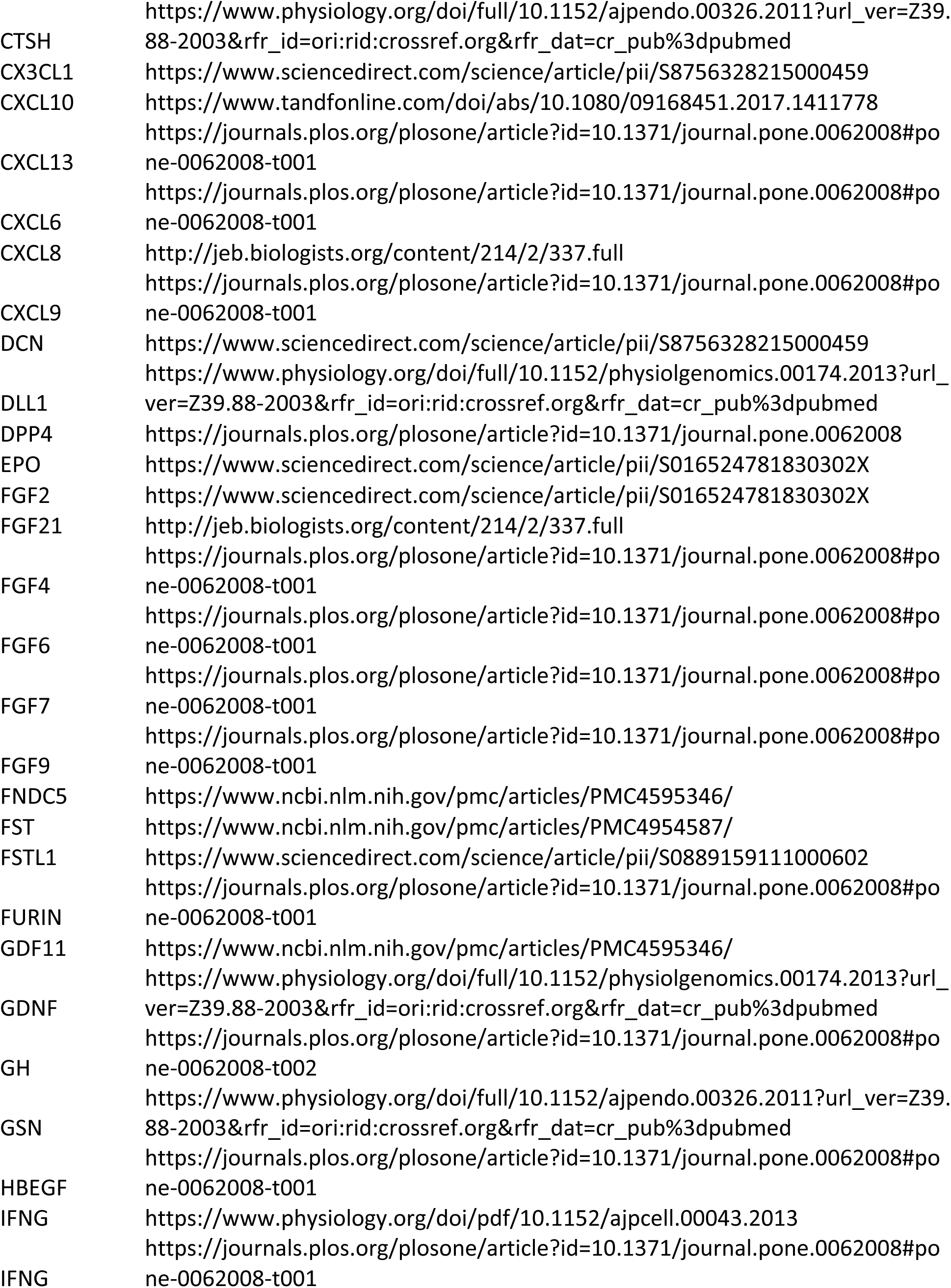
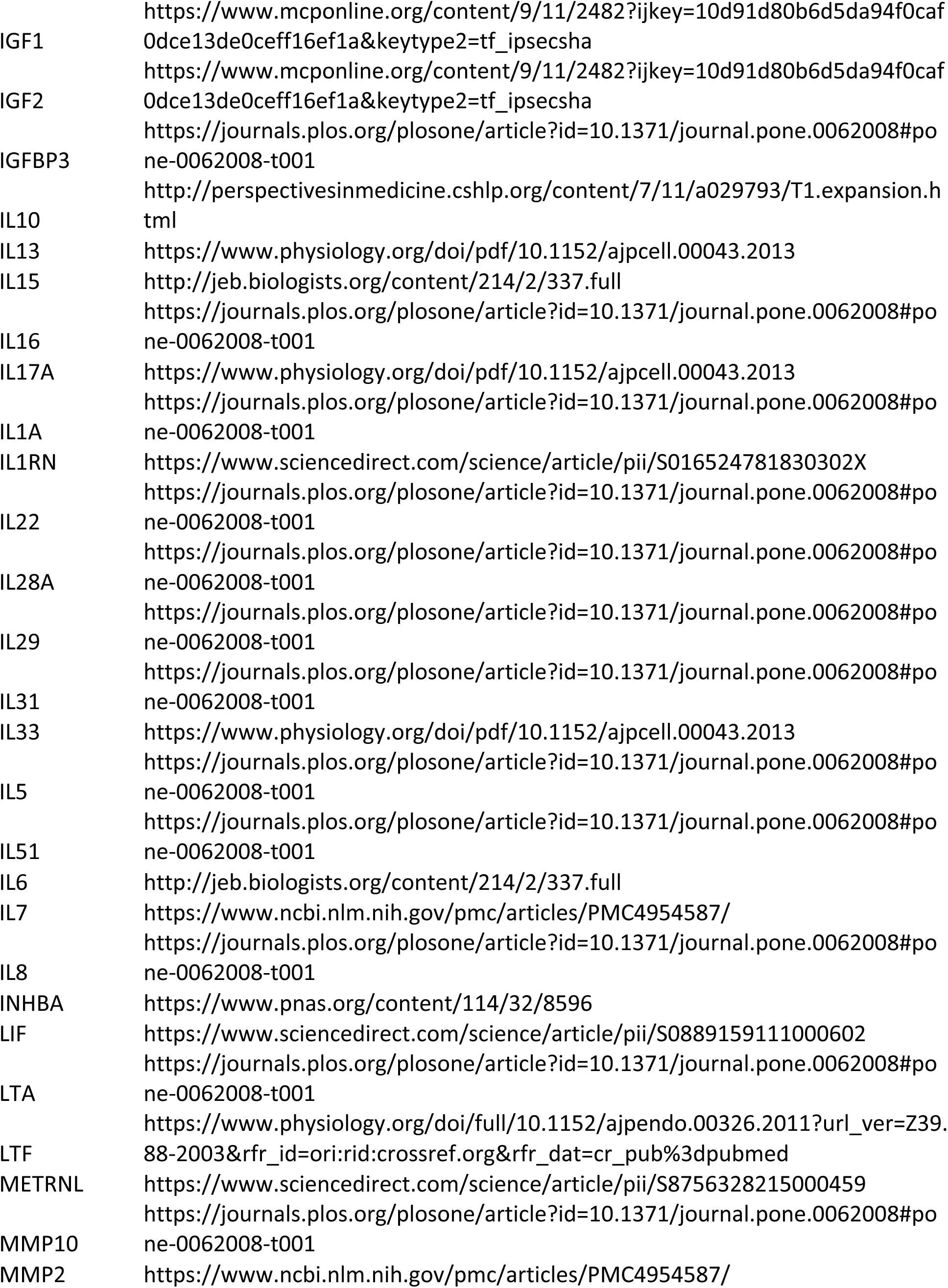
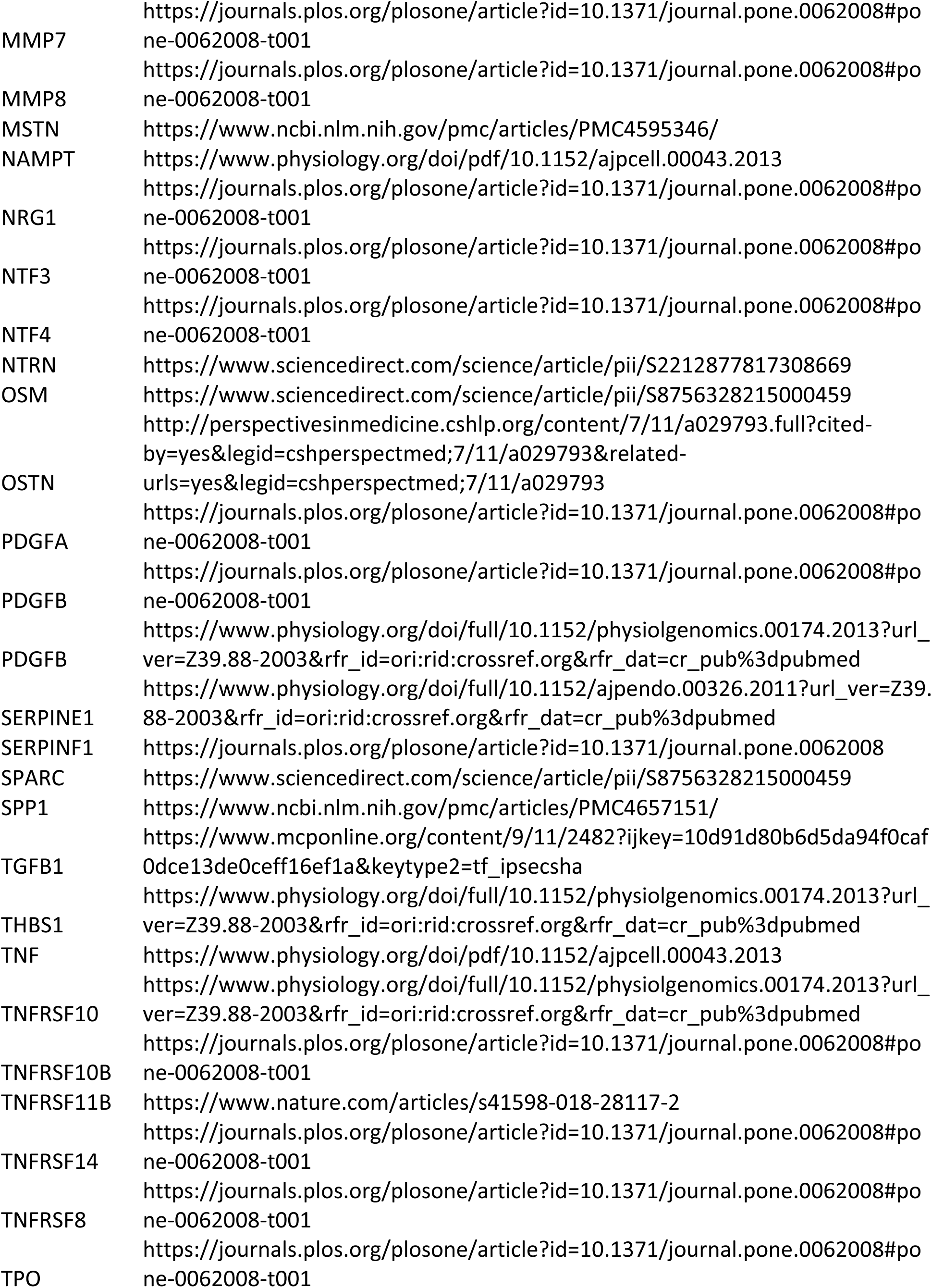
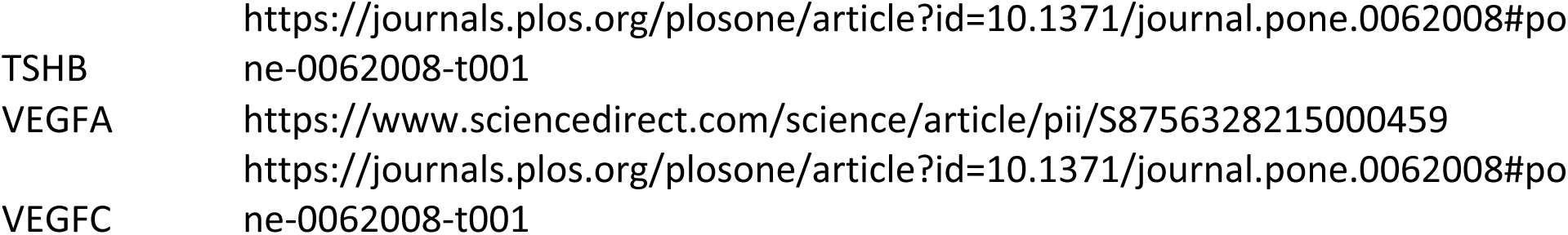
List of Myokines Based on Current Literature.

**Table S.3.**
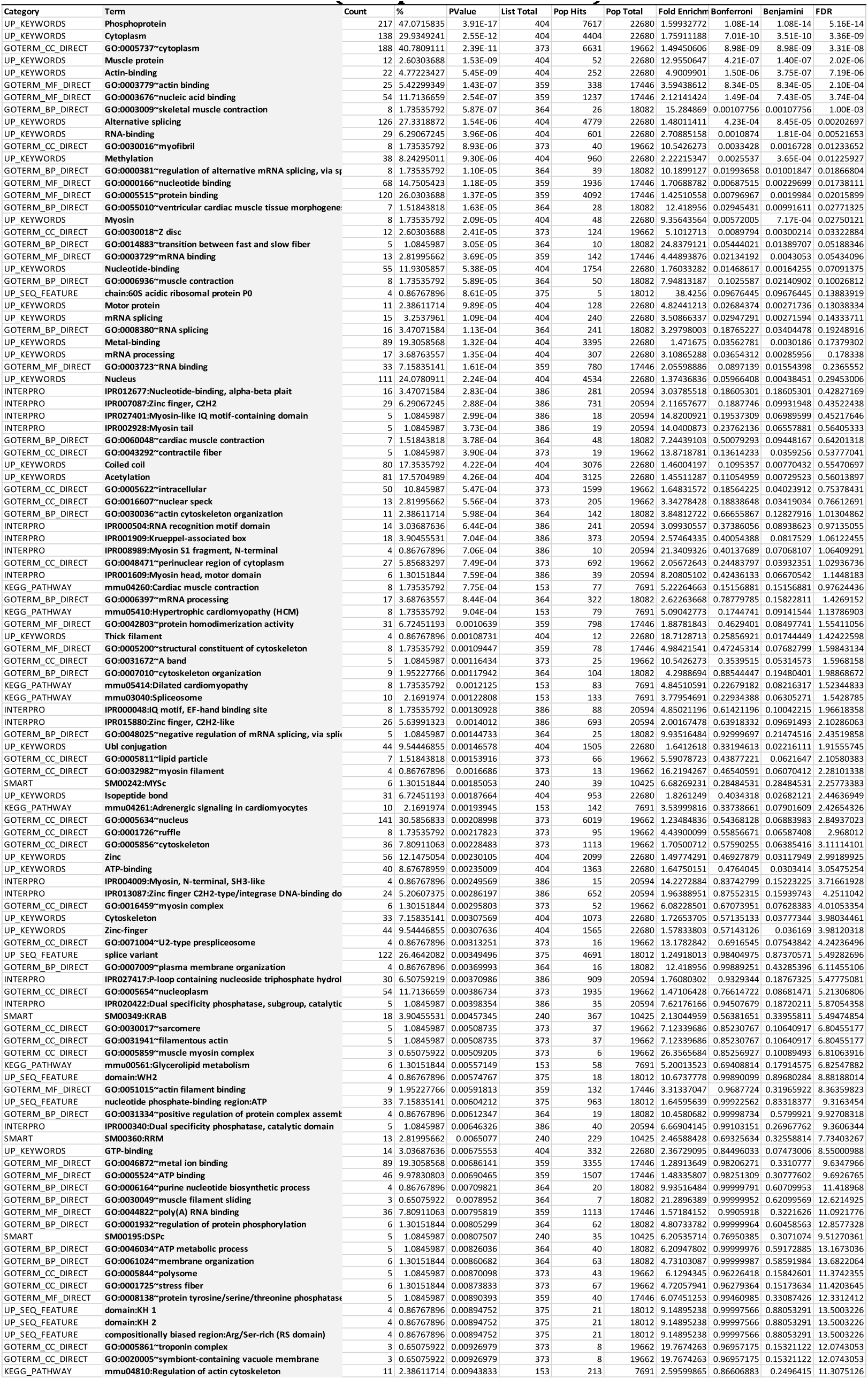
Gene Ontology (GO) analysis of genes up/downregulated in Myf6-knockout skeletal muscles (raw p-values <0.001)

**Table S.4.**
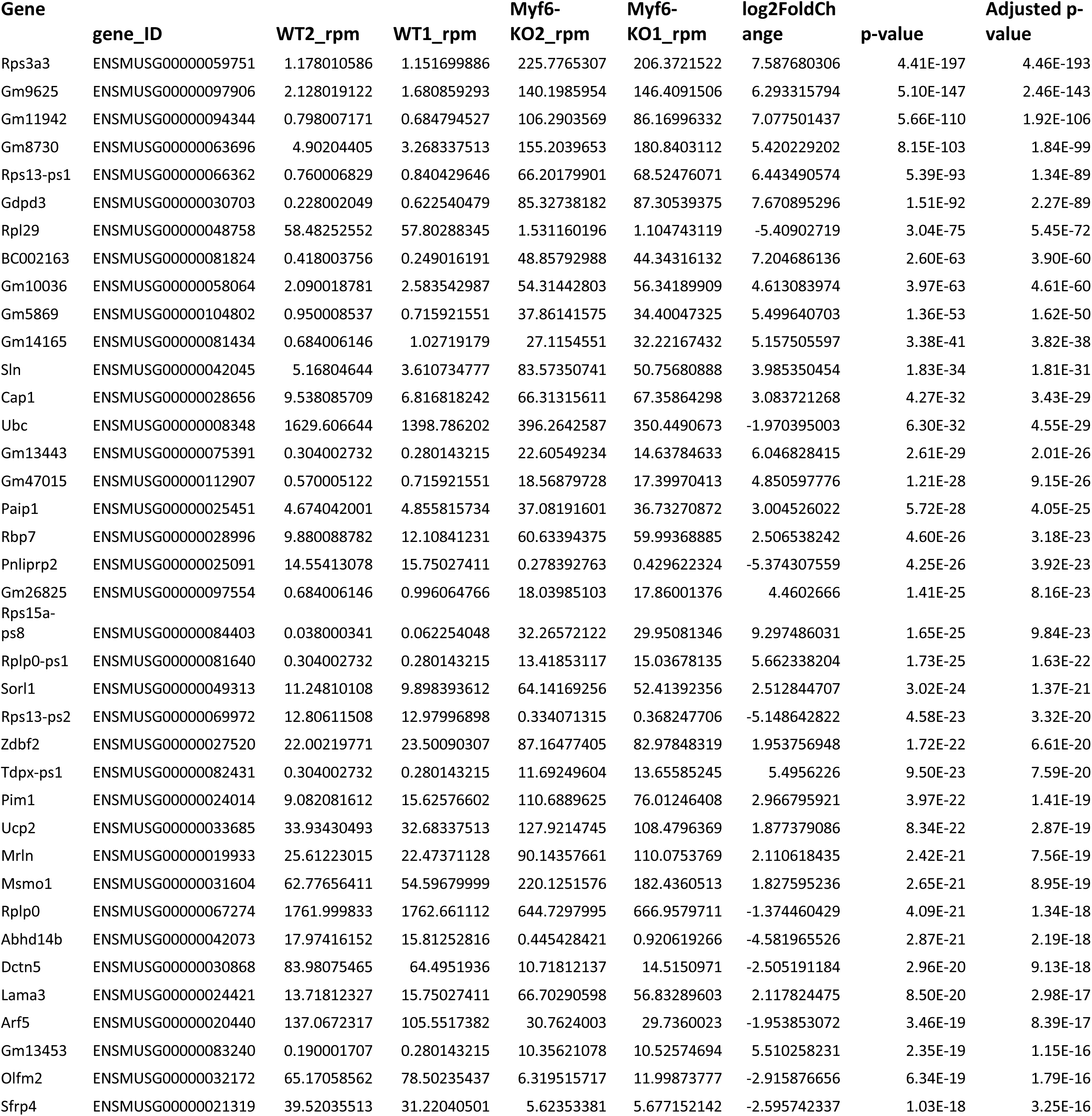
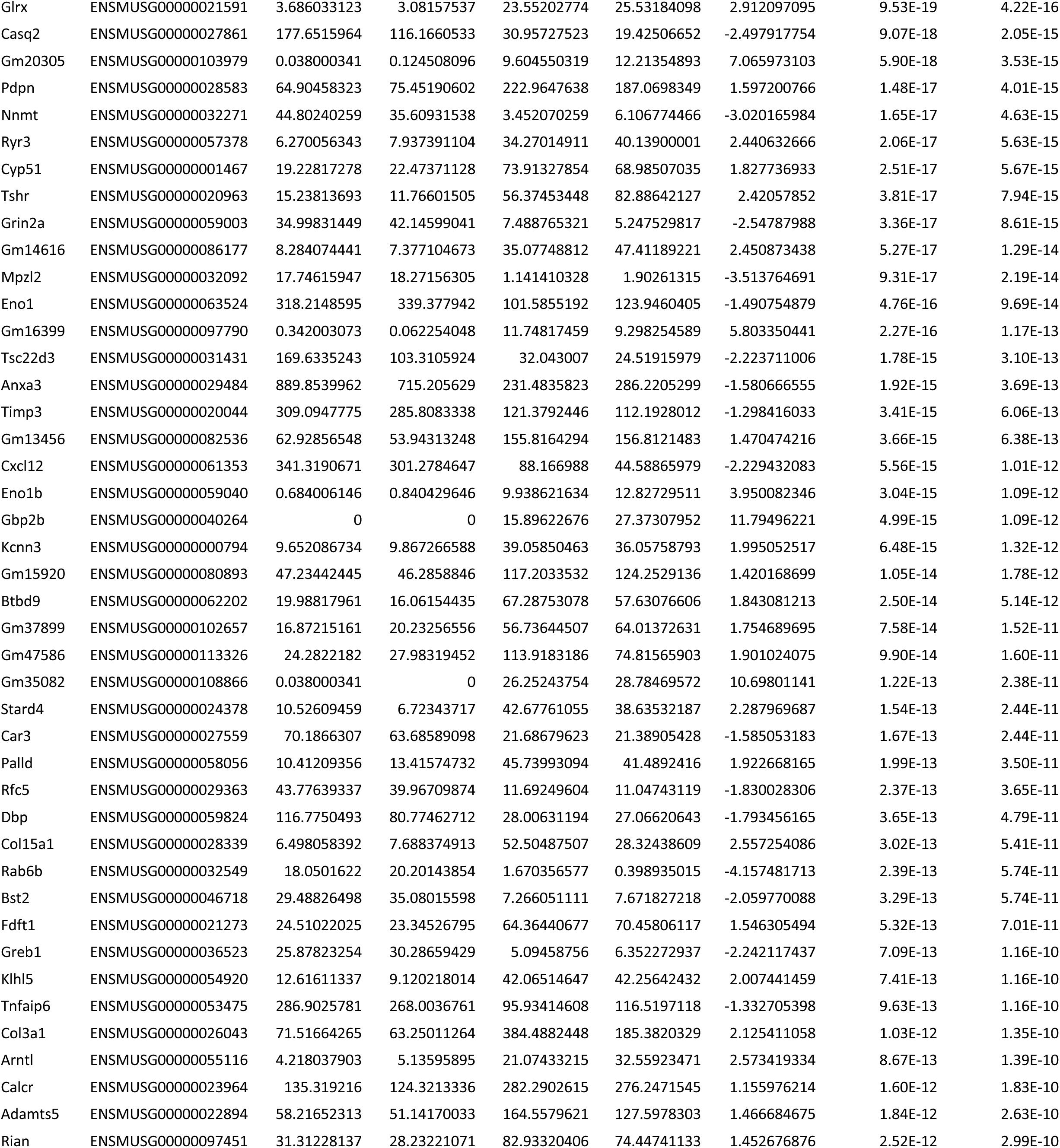
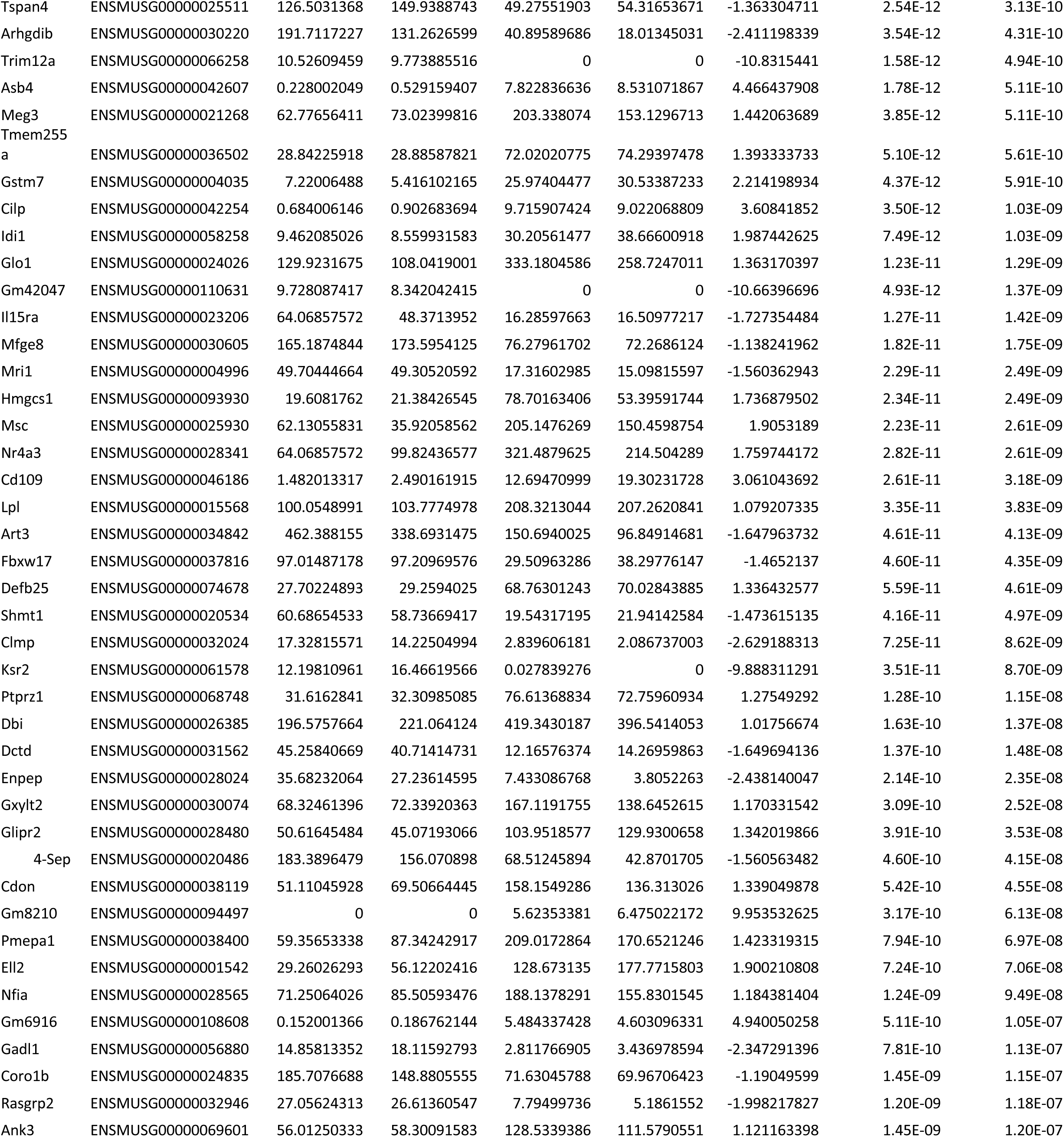
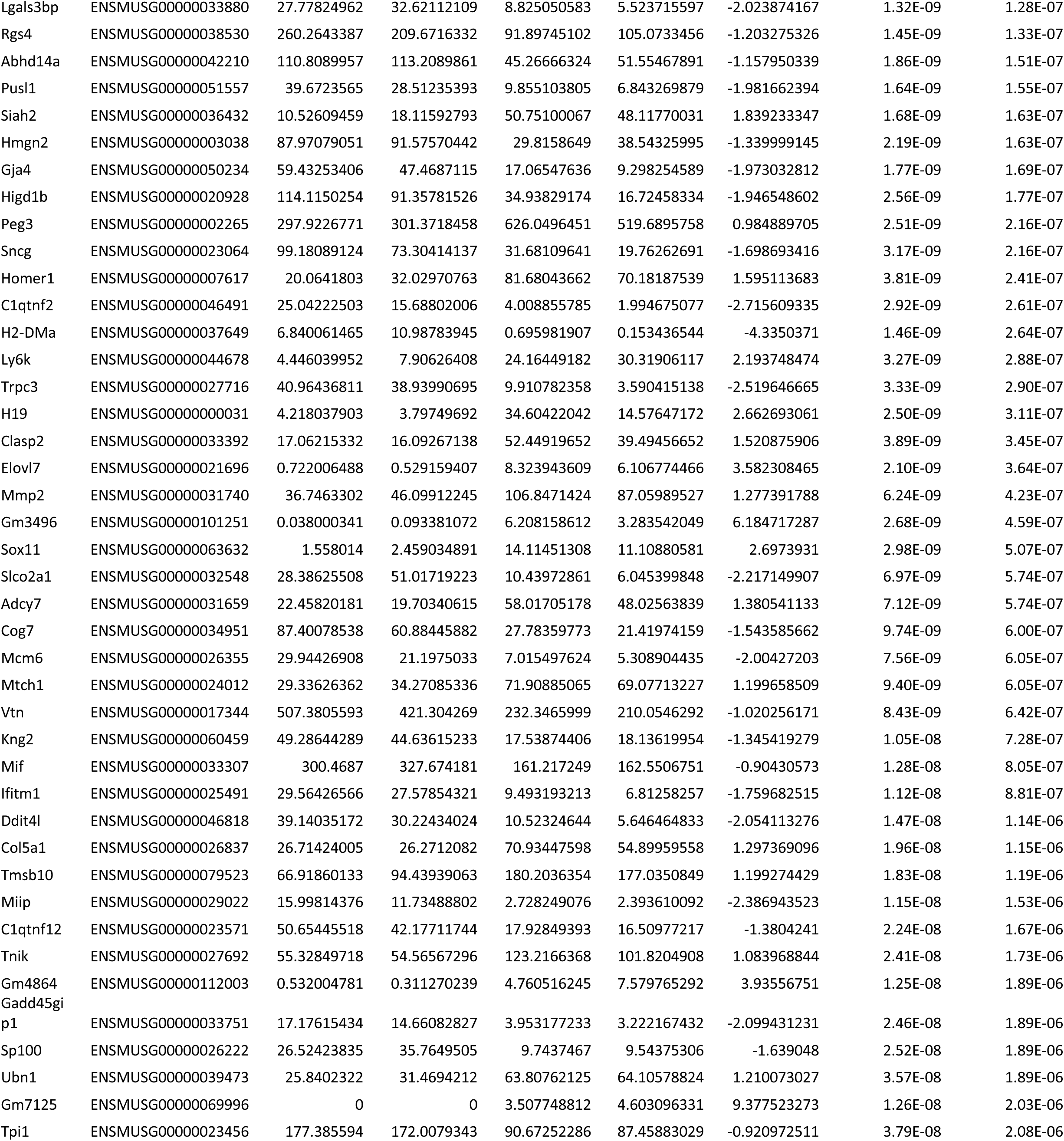
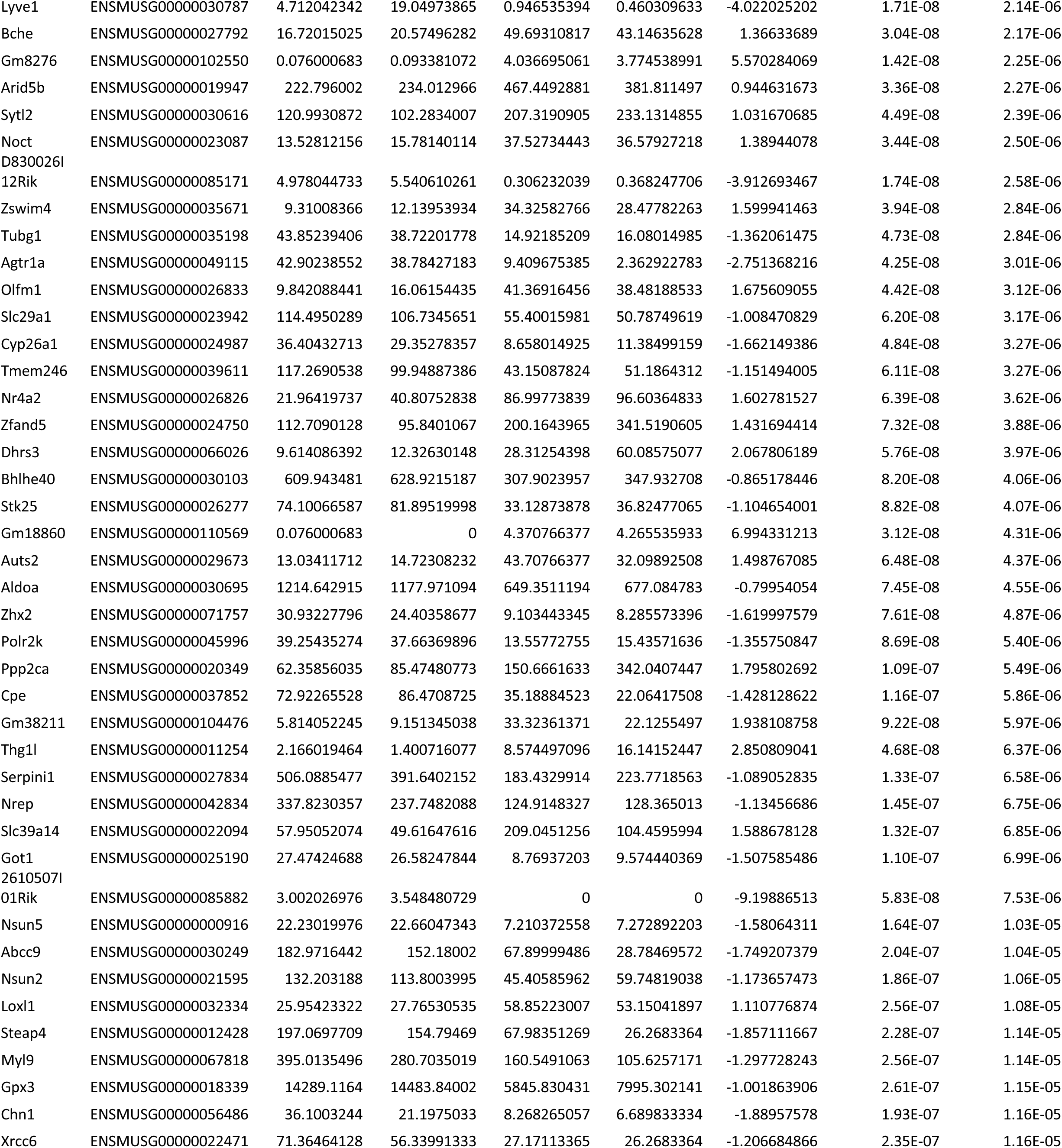
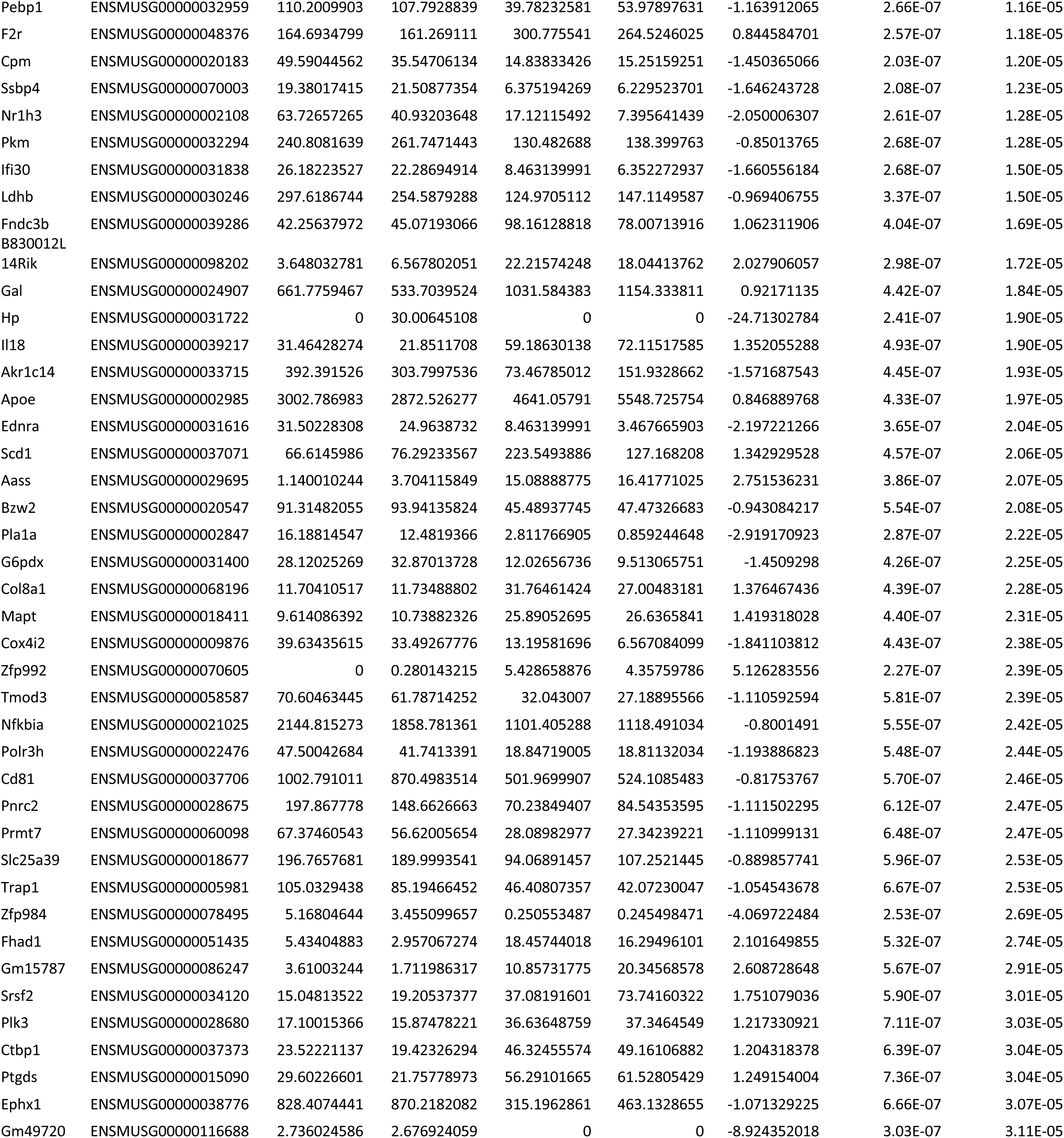
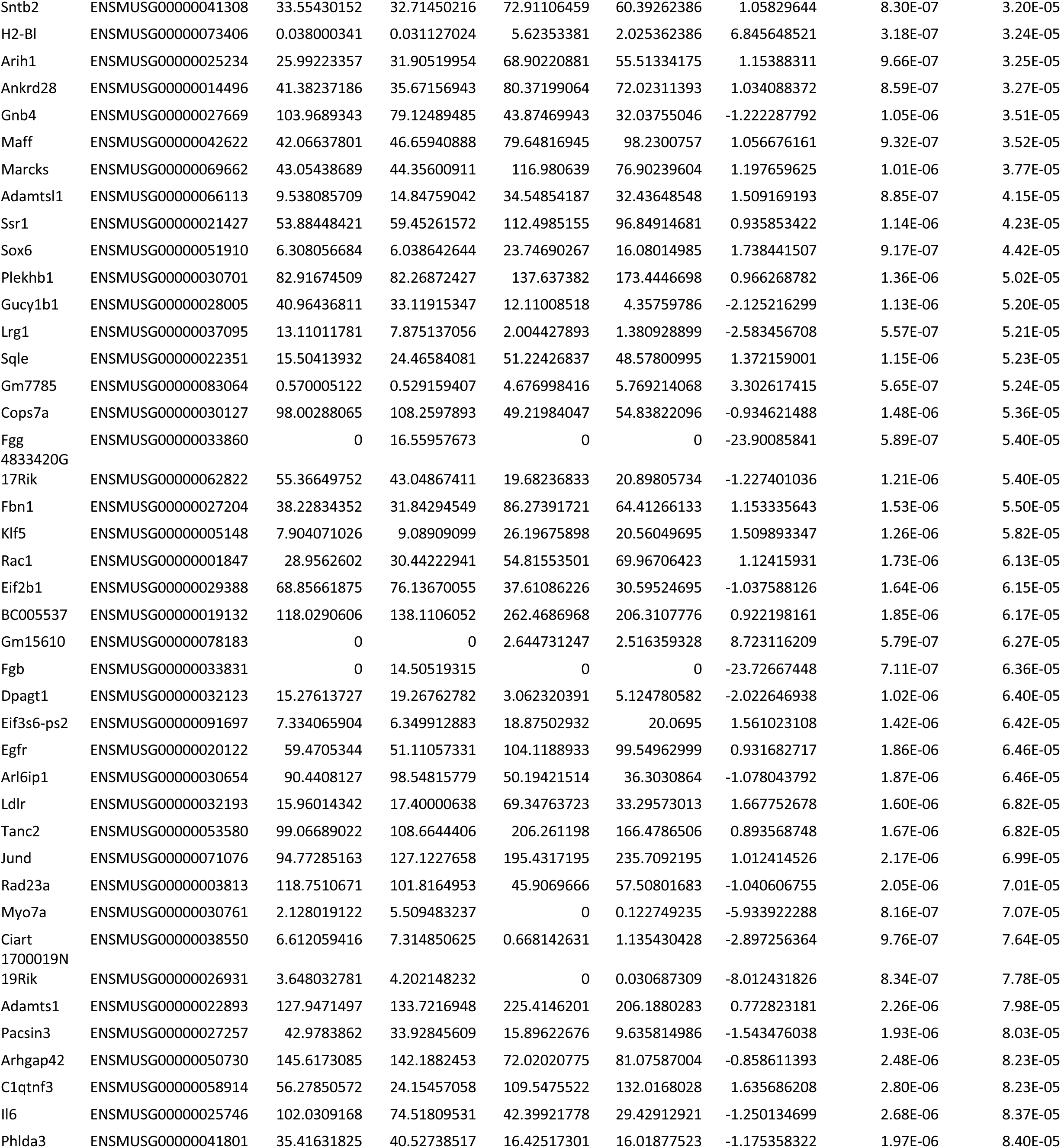
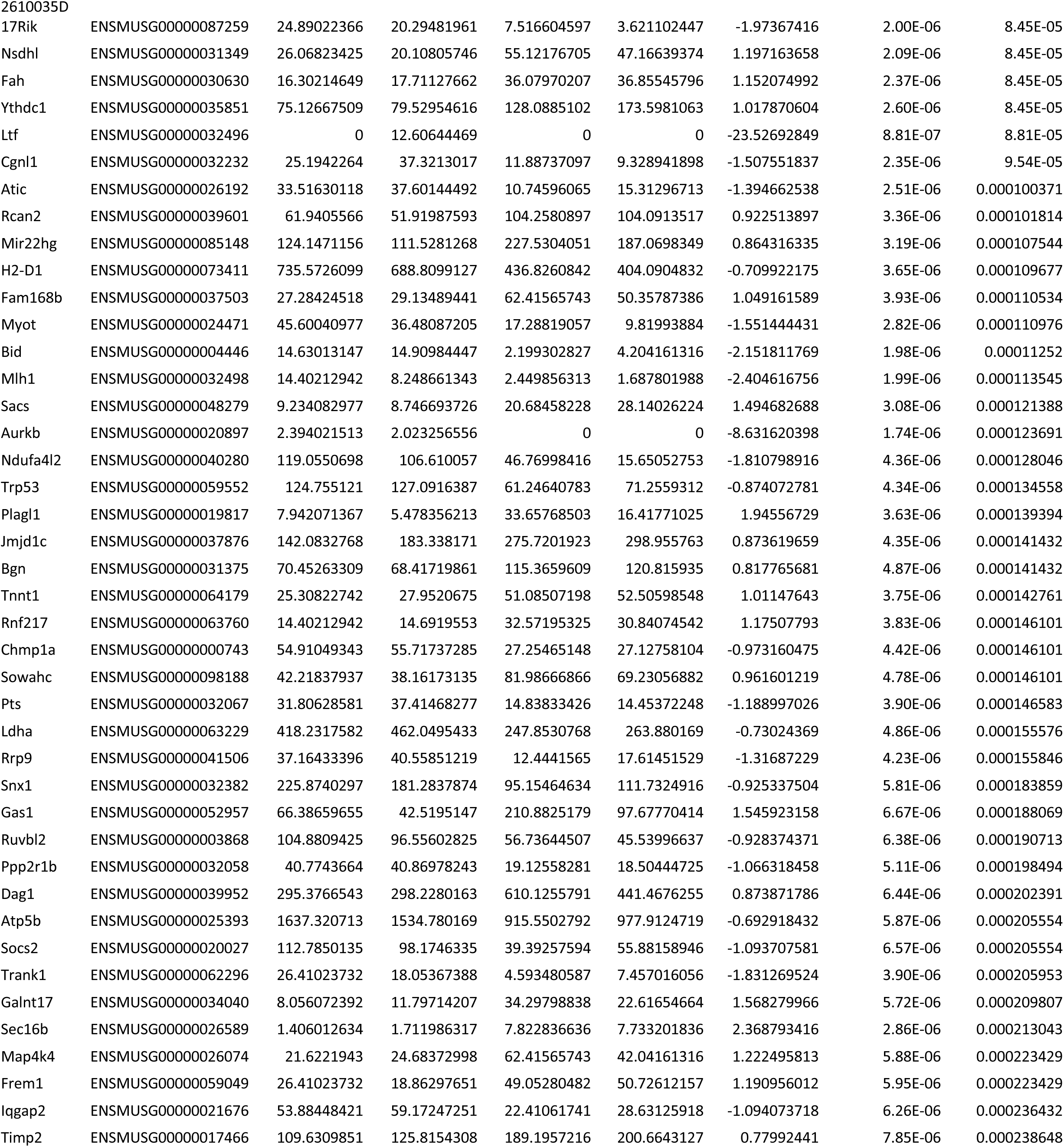
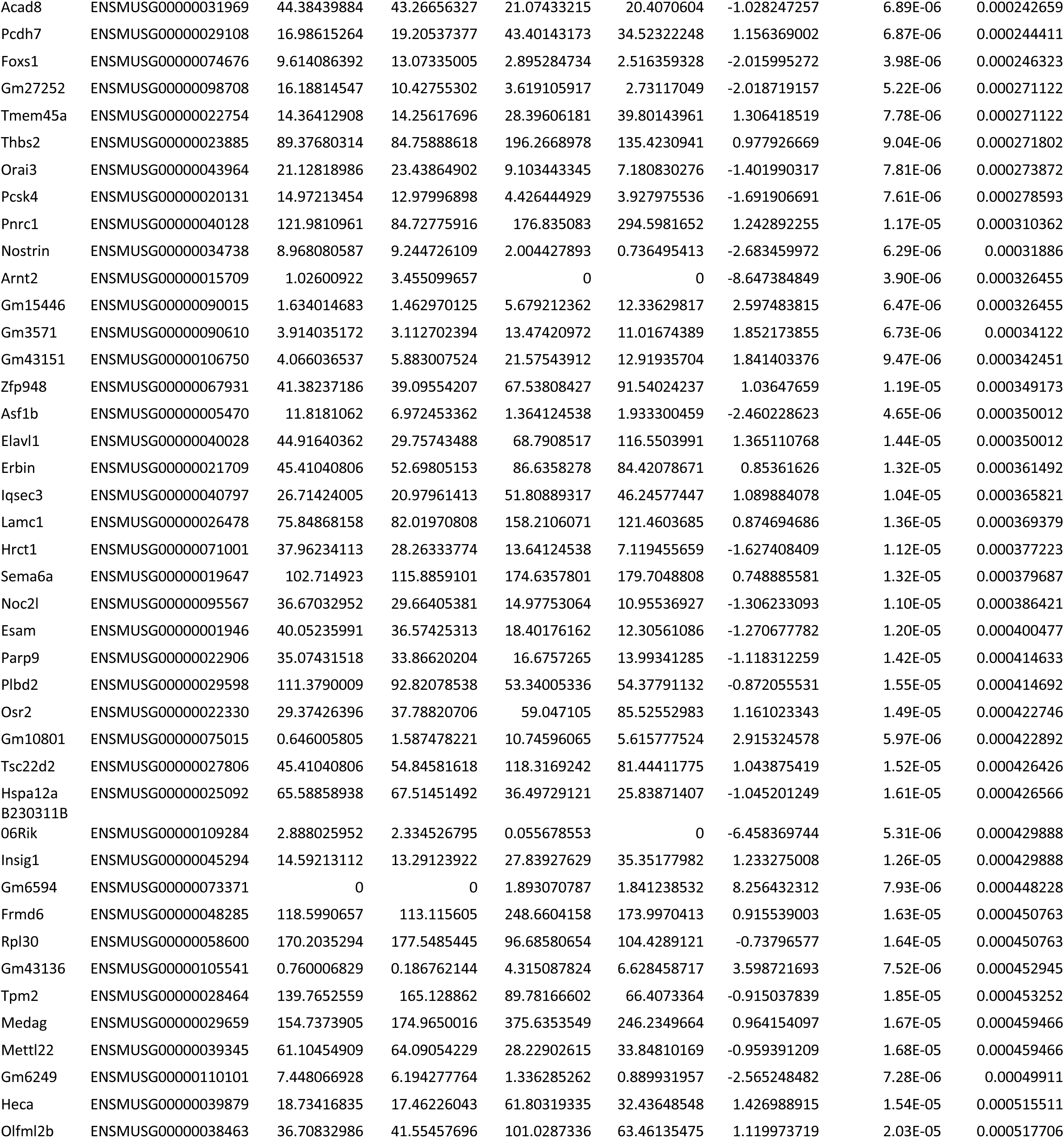
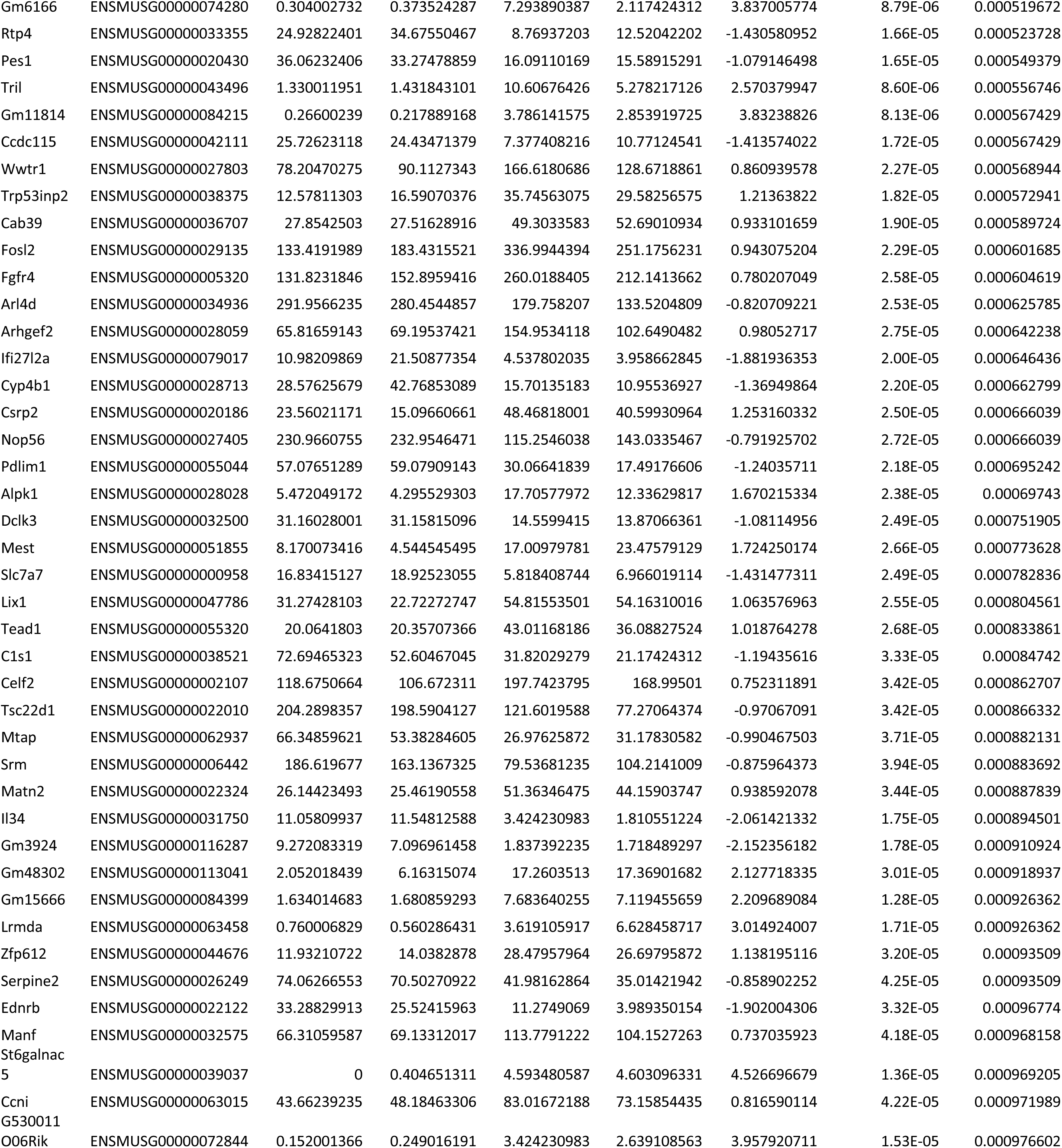
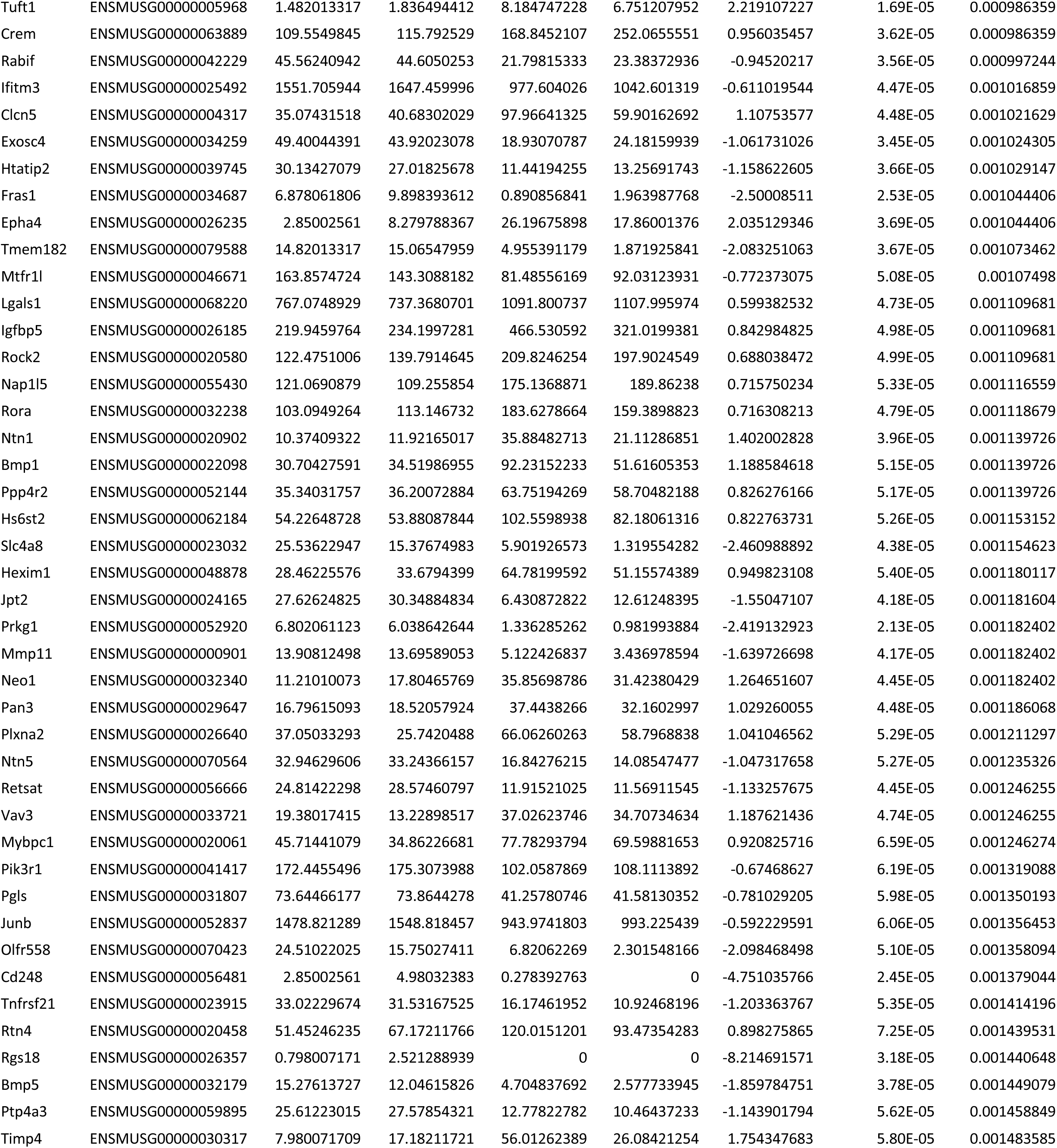
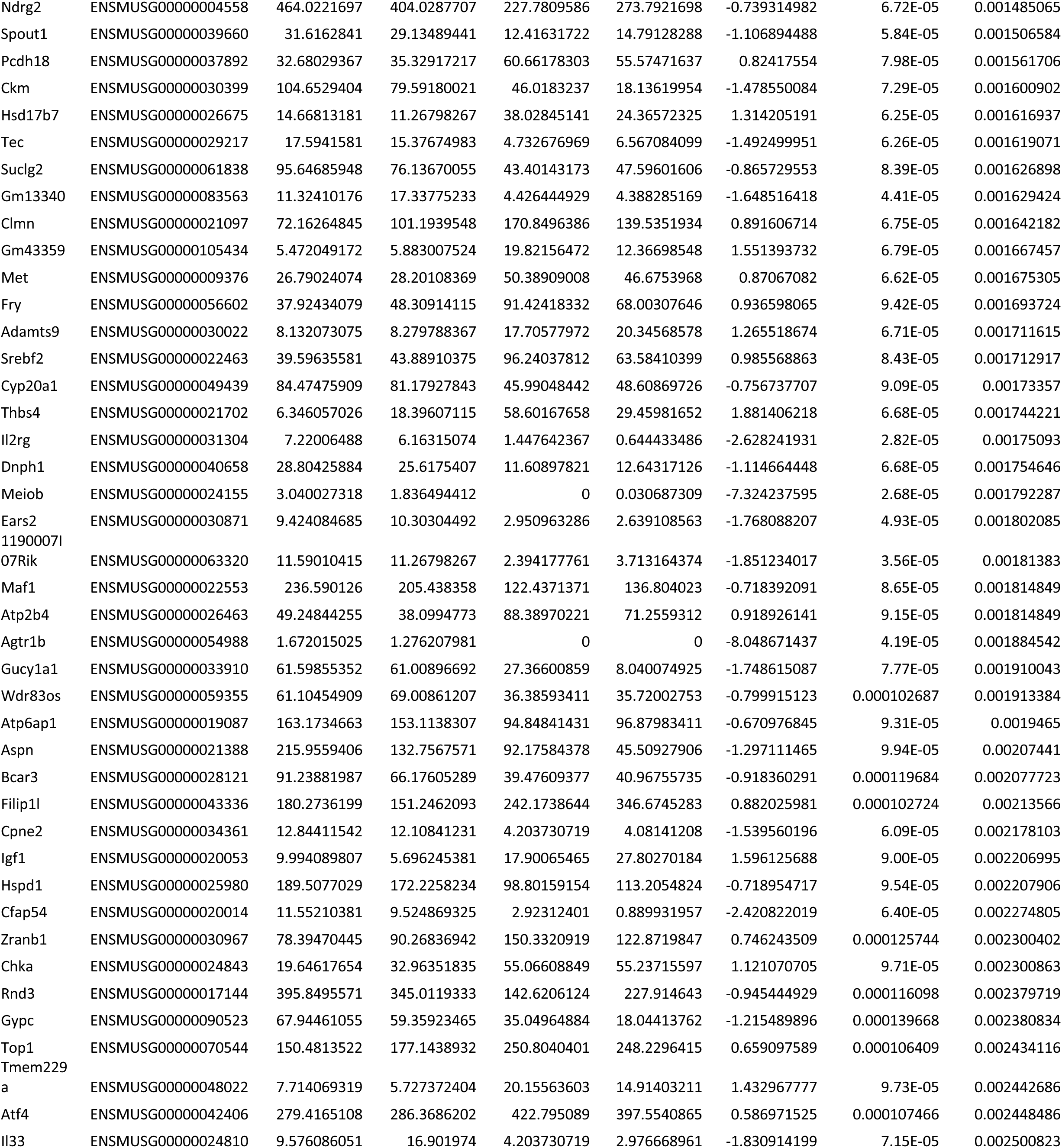
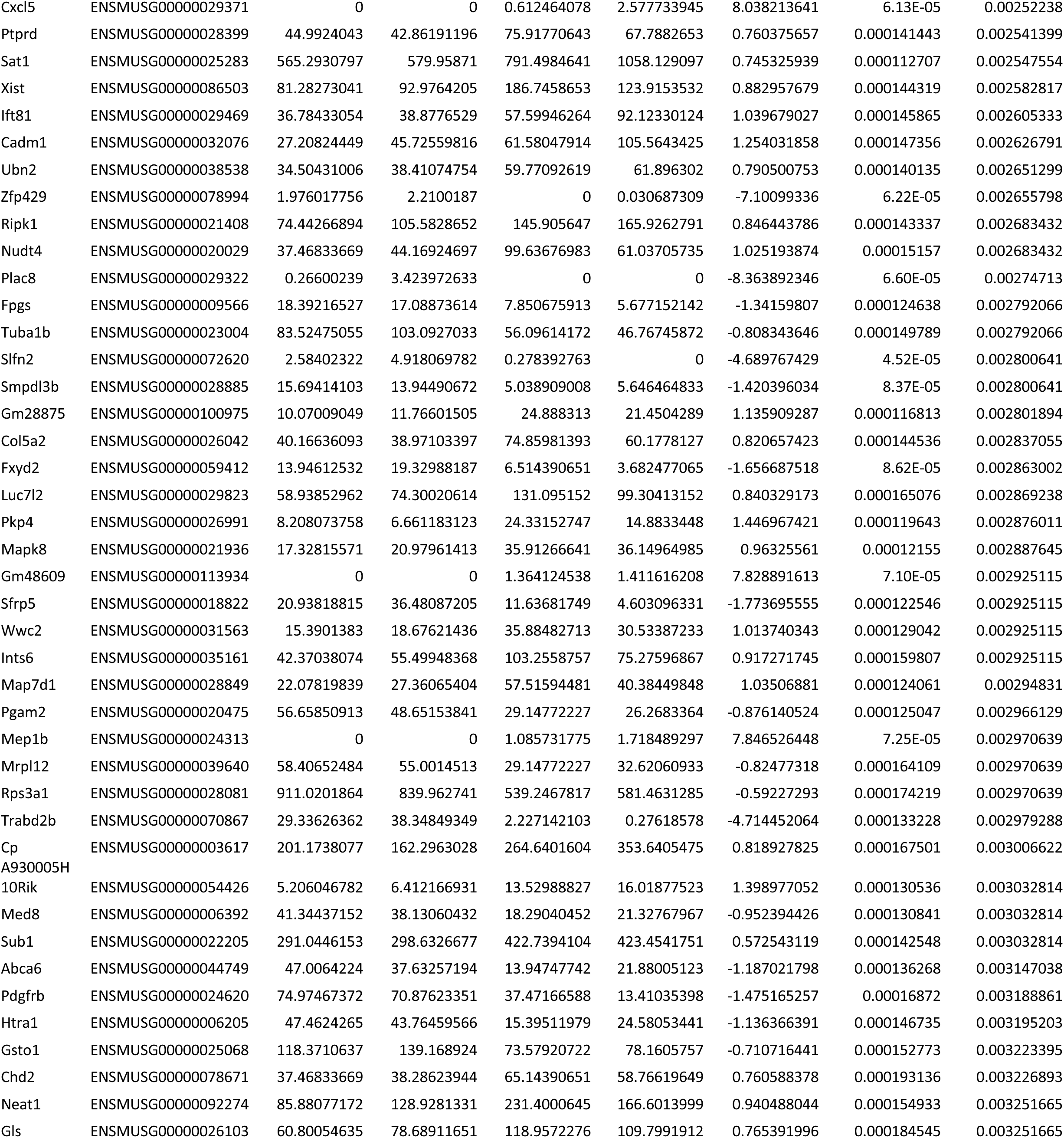
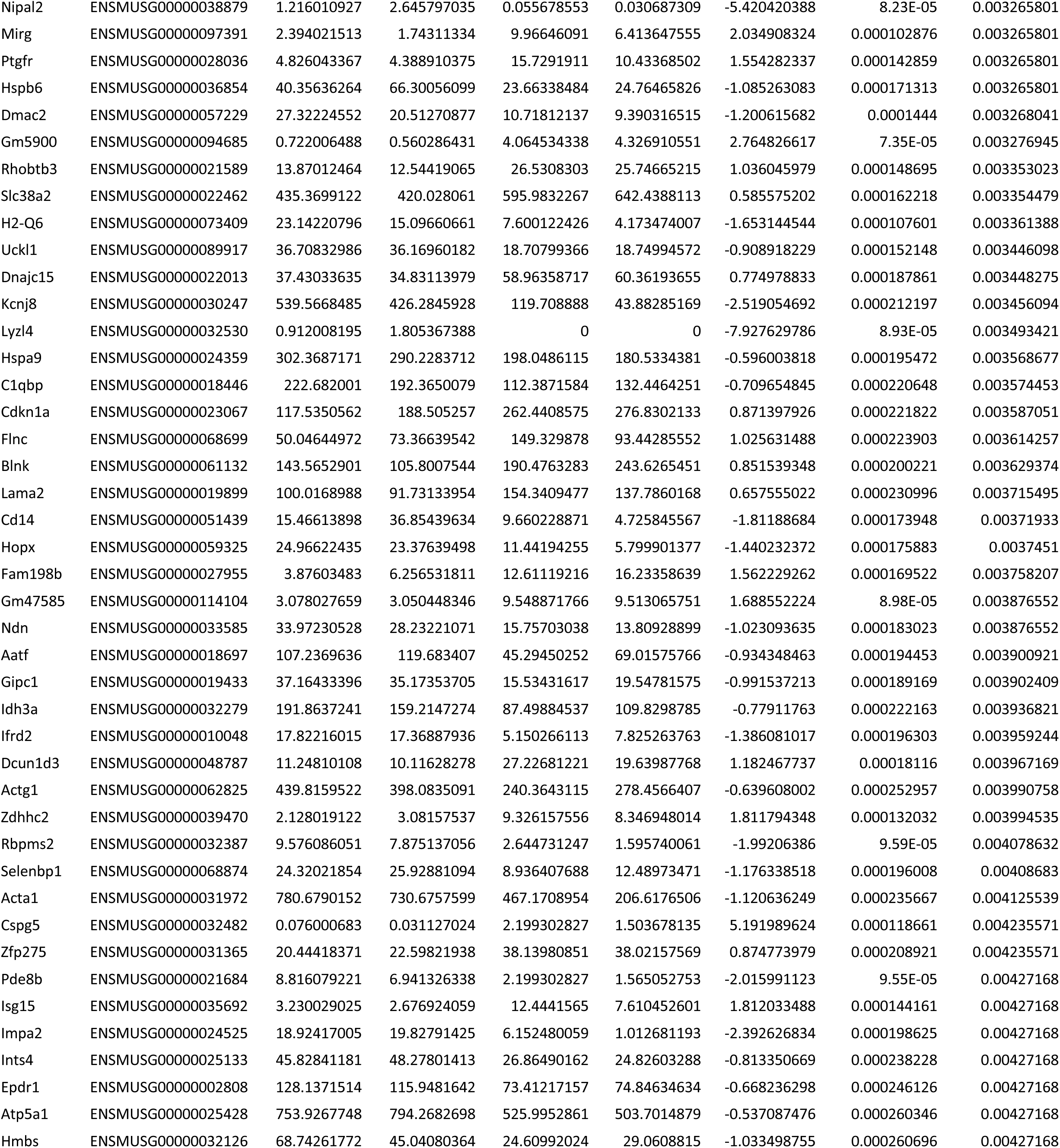
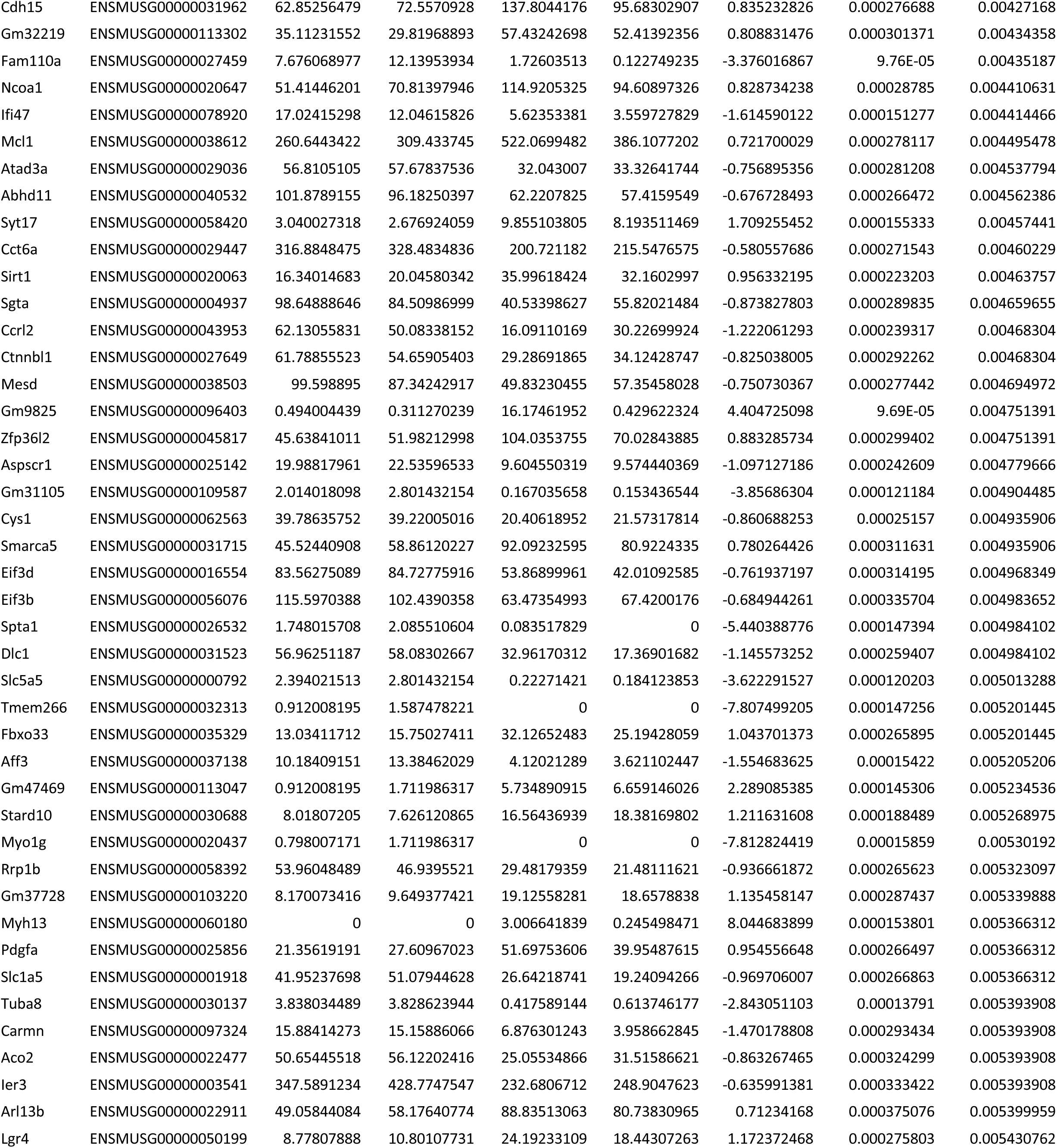
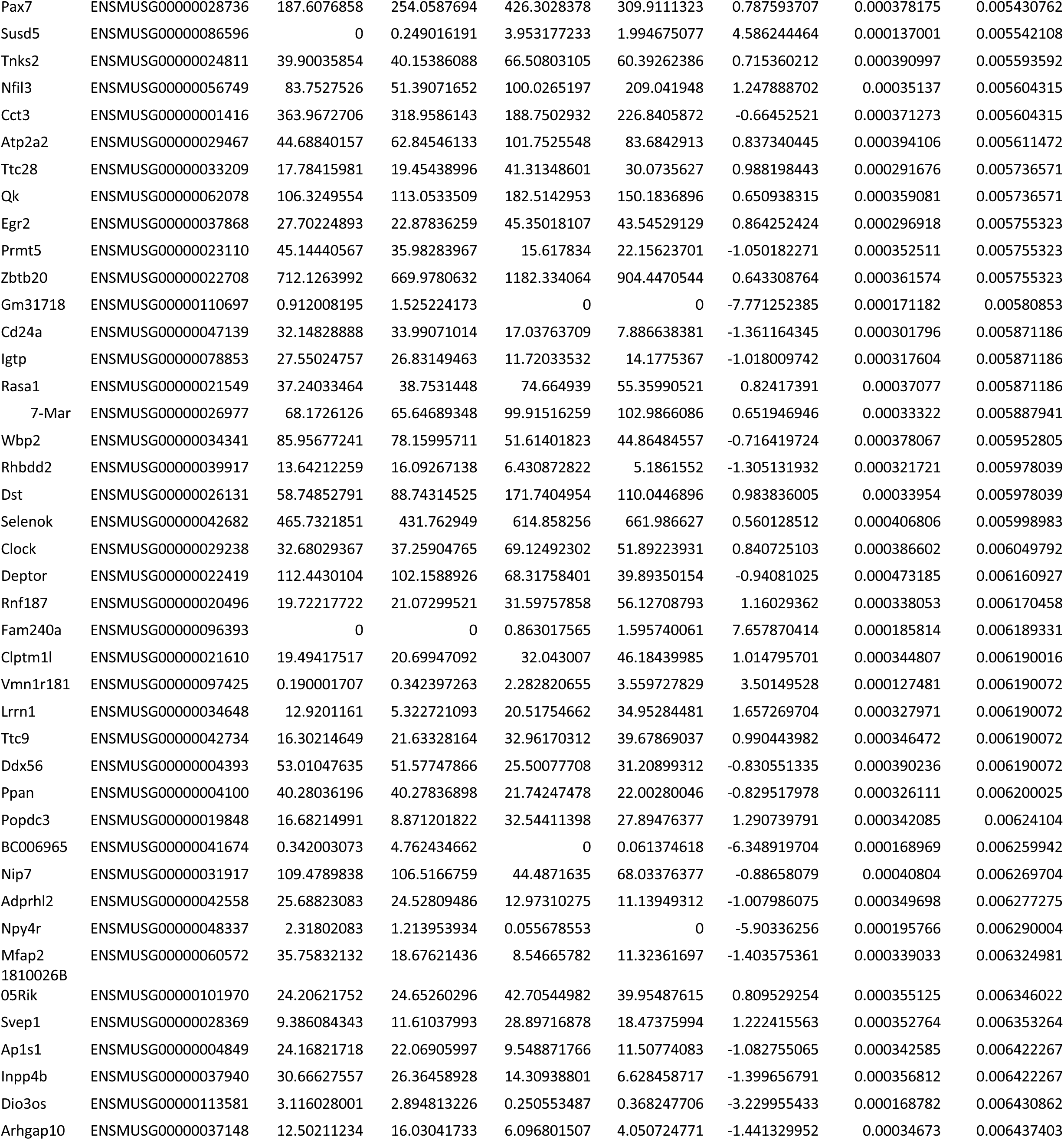
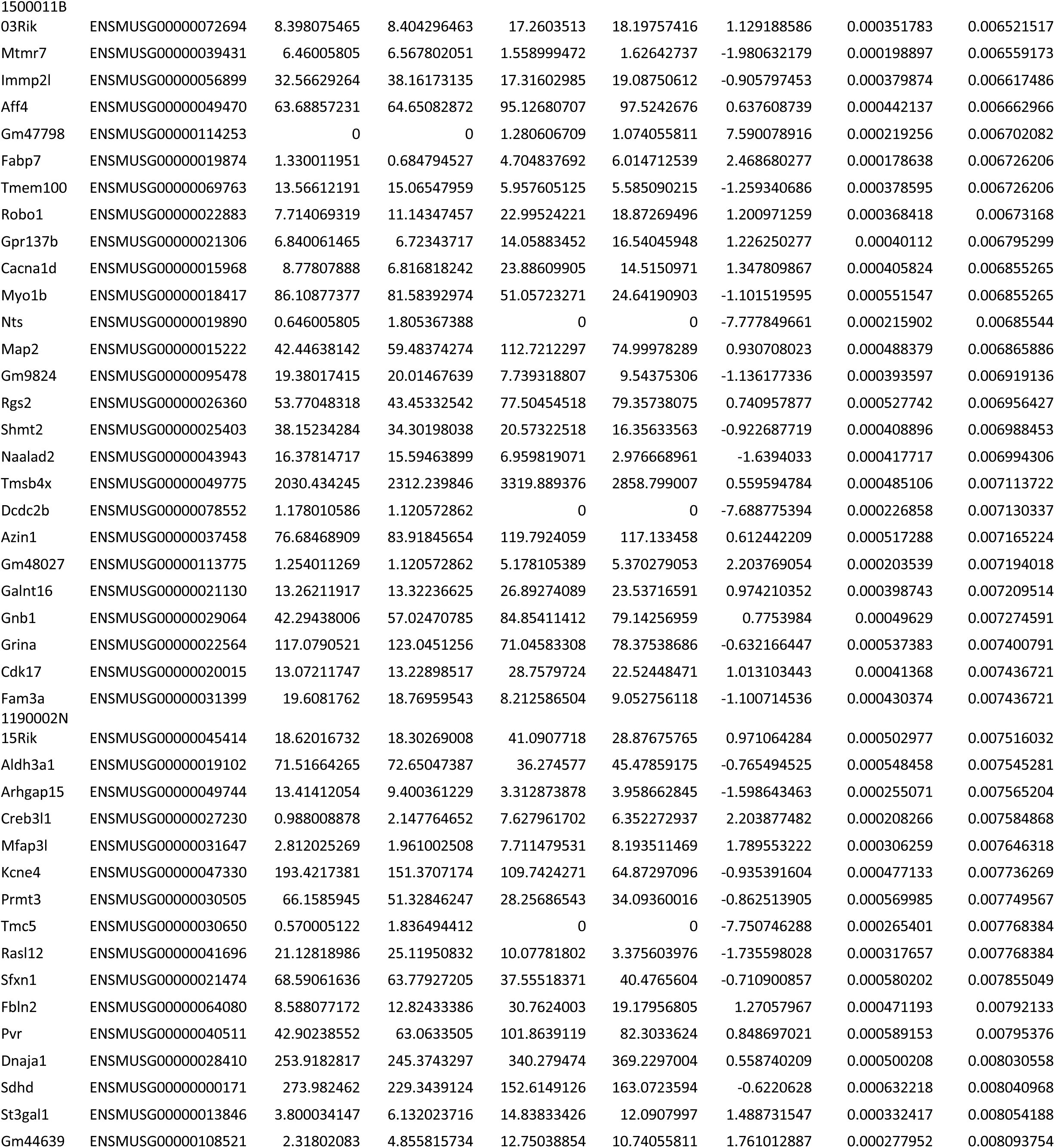
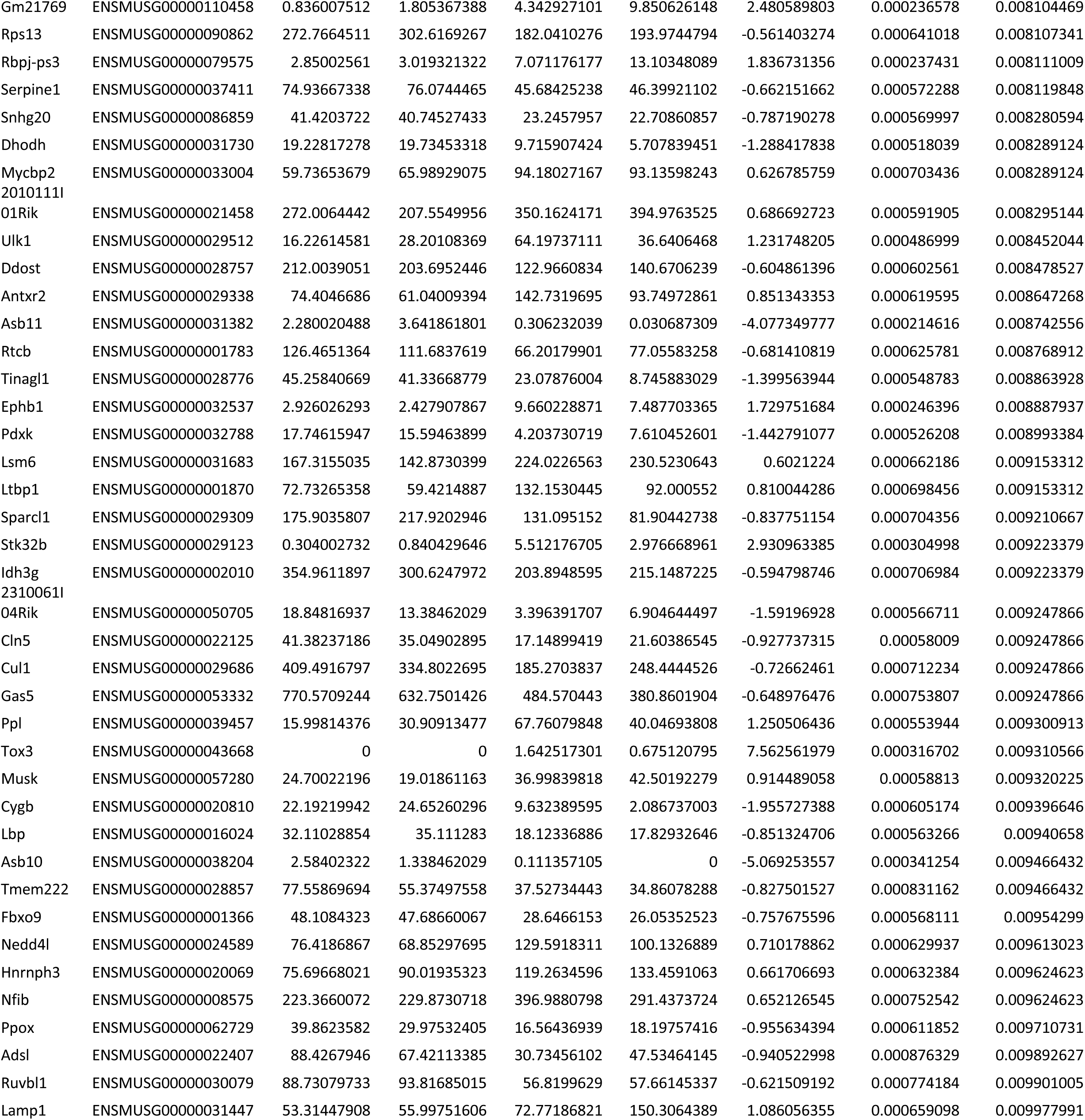
List of genes up/downregulated in Myf6-knockout satellite cells (adjusted p-values <0.01)

## Supplemental Figure Captions

**Figure S1.**
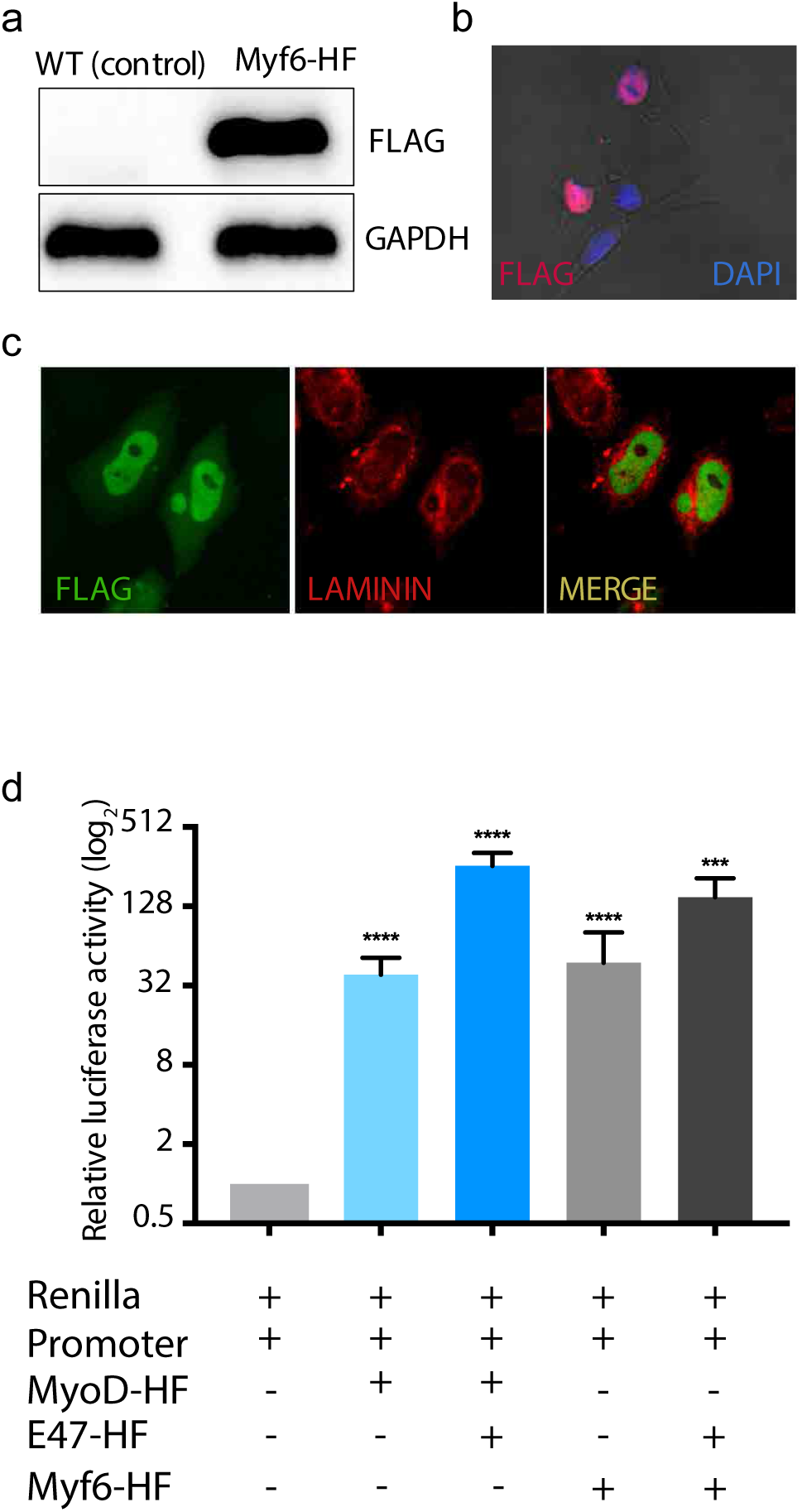
Functional Validation Myf6-CTAP Construct Used for ChIP-Seq Analysis. **(a)** Western Blot analysis of myoblasts infected with a retrovirus containing Myf6-CTAP. **(b)** Immunofluorescent analysis of myoblasts infected with a retrovirus containing Myf6-CTAP showing nuclear localization of the recombinant MYF6 protein using an antibody against FLAG (Monoclonal ANTI-FLAG M2 antibody, Sigma). **(c)** Transfection of Hela cells with Myf6-CTAP construct shows exclusive nuclear localization of the ectopically expressed MYF6-CTAP in Hela cells. **(d)** Dual luciferase assays showing the transcriptional activity of MYF6-CTAP on a synthetic regulatory sequence containing a trimeric canonical E-box motif (CAGCTG) spread over a 72bp stretch of a synthetic DNA. The synthetic sequence was sub cloned into the pGL4.23[luc2/minP] luciferase vector (Promega). Dual luciferase assays with Renilla/luciferase constructs were performed in Cos7 cells and a vector containing the full-length mouse MyoD (Soleimani et al., 2012a) was used as a positive control. Relative transcriptional activity of the MyoD and the Myf6 constructs were measured with and without co-transfection of Cos7 cells with the transcription factor E2-alpha (E47), a heterodimer of the myogenic regulatory factors (MRFs).

**Figure S2.**
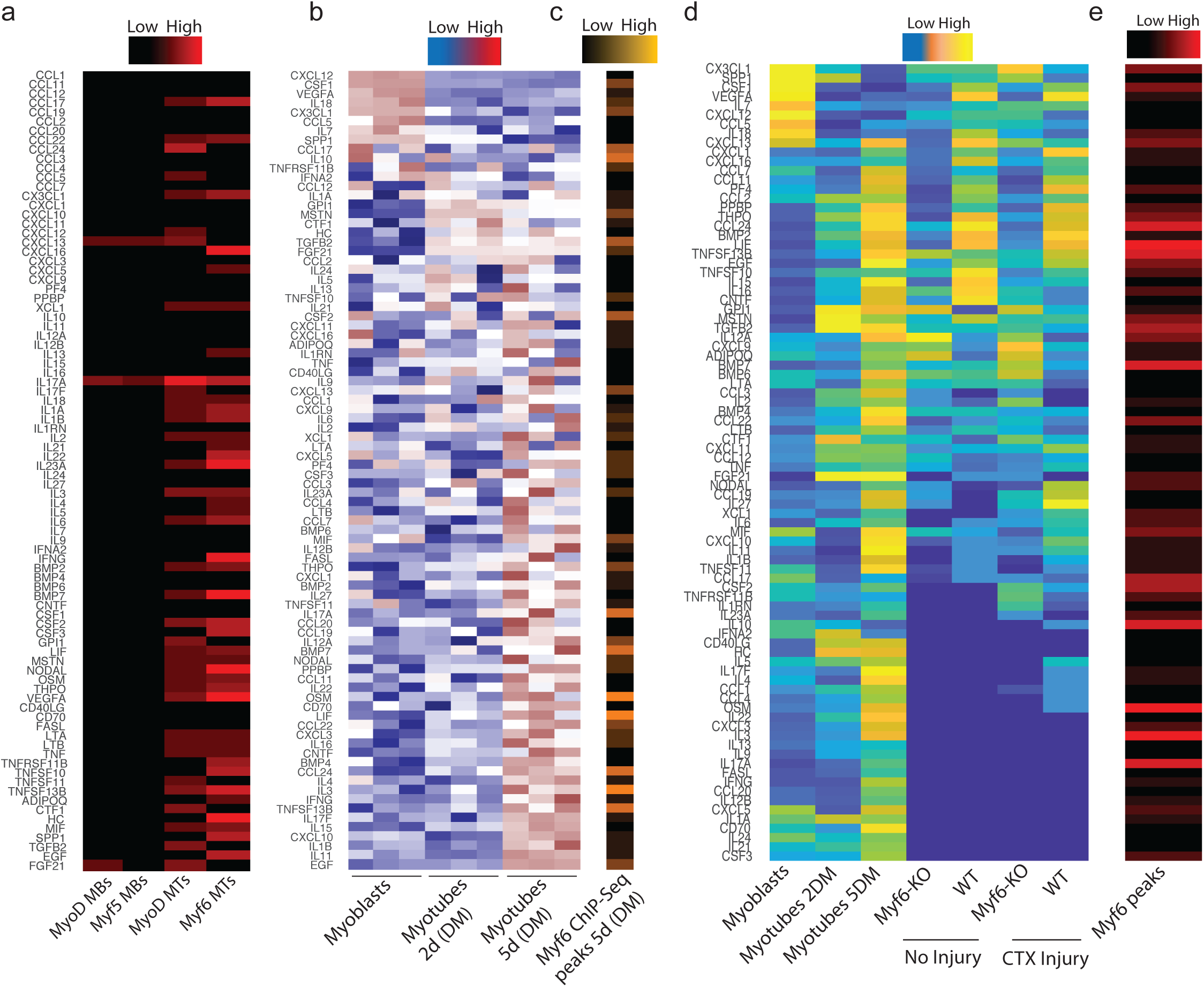
Myf6 Regulates the Expression of Various Cytokine Genes. **(a)** Colormap of Myf5, MyoD and Myf6 peaks within 100kb of the Transcription Start Sites (TSS) of cytokines ranging from zero (black) to six peaks (red) occupancy. Black indicates no binding (i.e., zero peaks), red indicates up to six ChIP-Seq peaks. The onset of differentiation coincides with increased binding of MRFs to the regulatory domains of the cytokines genes. **(b)** Gene expression analysis of cytokines during a five day time course of myogenic differentiation going from cycling myoblasts in growth media (Ham’s F10 supplemented with 20% Fetal Bovine Serum, 1% penicillin/streptomycin, 2.5 ng/ml basic Fibroblast Growth Factor) to terminally differentiated myocytes (2 days in differentiation media, DMEM supplemented with 5% horse serum) to the post mitotic multinucleated myotubes (5 days in differentiation media). Gene expression was assayed in biological triplicate by microarray (Soleimani et al., 2018; Soleimani et al., 2012a). Expression of each gene was averaged across three replicates and normalized to mean zero and standard deviation of one. Expression values were then truncated at +/−3 for color display. Blue indicates lower than average expression, while red indicates higher than average expression. **(c)** Distribution of Myf6 peaks within 100kb of the TSS of the cytokines genes, similar to the analysis shown in “a”. **(d)** Color map of genes that are up/down regulated in the whole muscle transcriptome (RNA-Seq) of hindlimb muscles from Myf6-KO and WT counterparts. Differential expression is Log_2_ transformed and truncated at +/−3. Color scale goes from blue (−3=lower than mean), to white (0=mean) and finally to red (+3=higher than mean).

**Figure S3.**
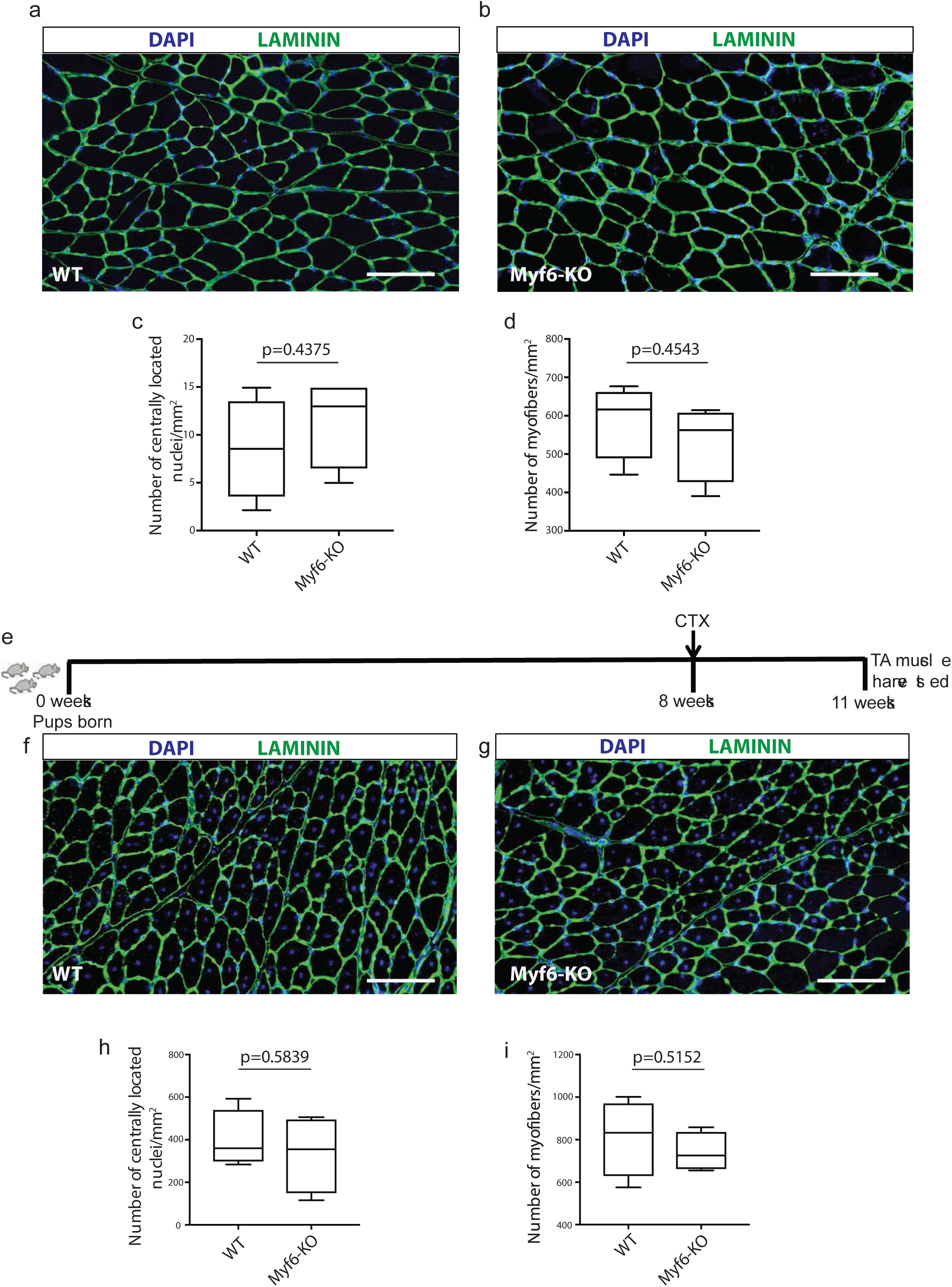
Myf6 is Dispensable for Muscle Development and Regeneration in Adult Mice. **(a-b)** Immunofluorescent analysis of TA muscles from WT and Myf6-KO counterparts stained with Laminin (LAMA1) and counterstained with DAPI. **(c)** Quantification of the number of regenerating fibers (depicted by the presence of centrally located myonuclei) between the WT and the Myf6-KO mice. **(d)** Quantification of the number of myofibers per unit area (mm2) of the TA muscles between WT and Myf6-KO counterparts. **(e)** Immunofluorescent analysis of TA muscle from WT and Myf6-KO animals three weeks after CTX injury (see extended Materials and Methods) stained with Laminin and counterstained with DAPI. **(f)** Quantification of the number of regenerating myofibers (centrally located myonuclei) between WT and Myf6-KO counterparts. **(g)** Quantification of the number of myofibers per unit area (mm2) between WT and Myf6-KO counterparts. (n=4 animals per group were used. p values are based on two tailed t-test. Error bars represent standard deviation).

**Figure S4.**
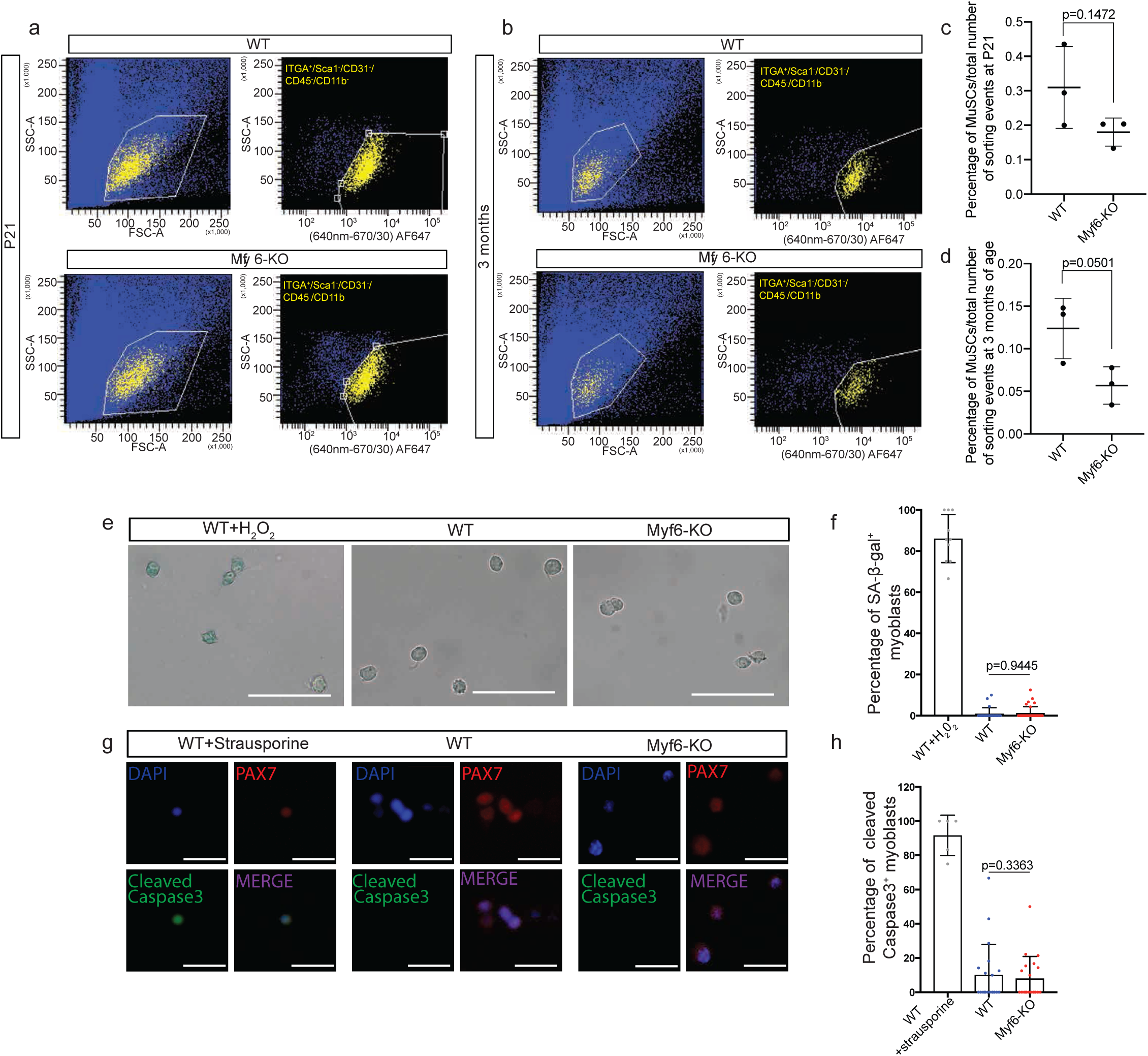
Myf6-Knockout Mice Show a Reduced MuSC Pool. **(a)** Representative FACS plots for the isolation of MuSCs from Myf6-KO and WT mice at P21. The yellow population corresponds to the ITGA7+/CD31-/SCA1-/CD45-/CD11b-population of MuSCs. **(b)** Quantification of the percentage of total FACS sorting events corresponding to the MuSC population (number of ITGA+/ ITGA7+/CD31-/SCA1-/CD45-/CD11b-events over the total number of FACS events) at P21. **(c)** Representative FACS plots for the isolation of MuSCs from Myf6-KO and WT mice at 3 months. The yellow population corresponds to the ITGA7+/CD31-/SCA1-/CD45-/CD11b-population of MuSCs. **(d)** Quantification of the percentage of total FACS sorting events corresponding to the MuSC population (number of ITGA+/ ITGA7+/CD31-/SCA1-/CD45-/CD11b-events over the total number of FACS events) at 3 months. **(e)** Representative photos of SA-β-galactosidase staining in Myf6-KO, WT and WT H_2_O_2_-treated cultured primary myoblasts. As a positive control, primary myoblasts from WT mice were cultured and treated with 20μM H_2_O_2_ for 15 minutes, and left in fresh media for 48 hours before fixation **(f)** Quantification of the percentage of SA-β-Gal+ cells in Myf6-KO, WT and WT H_2_O_2_-treated myoblasts. **(g)** Representative photos of staining for cleaved caspase 3 in Myf6-KO, WT, and WT strausporine-treated cultured primary myoblasts. Primary myoblasts from WT mice were cultured and treated with 3μM strausporine for 3 hours and then fixed. **(h)** Quantification of the percentage of cleaved caspase3+ cells in Myf6-KO, WT and WT strausporine-treated myoblasts.

**Figure S5.**
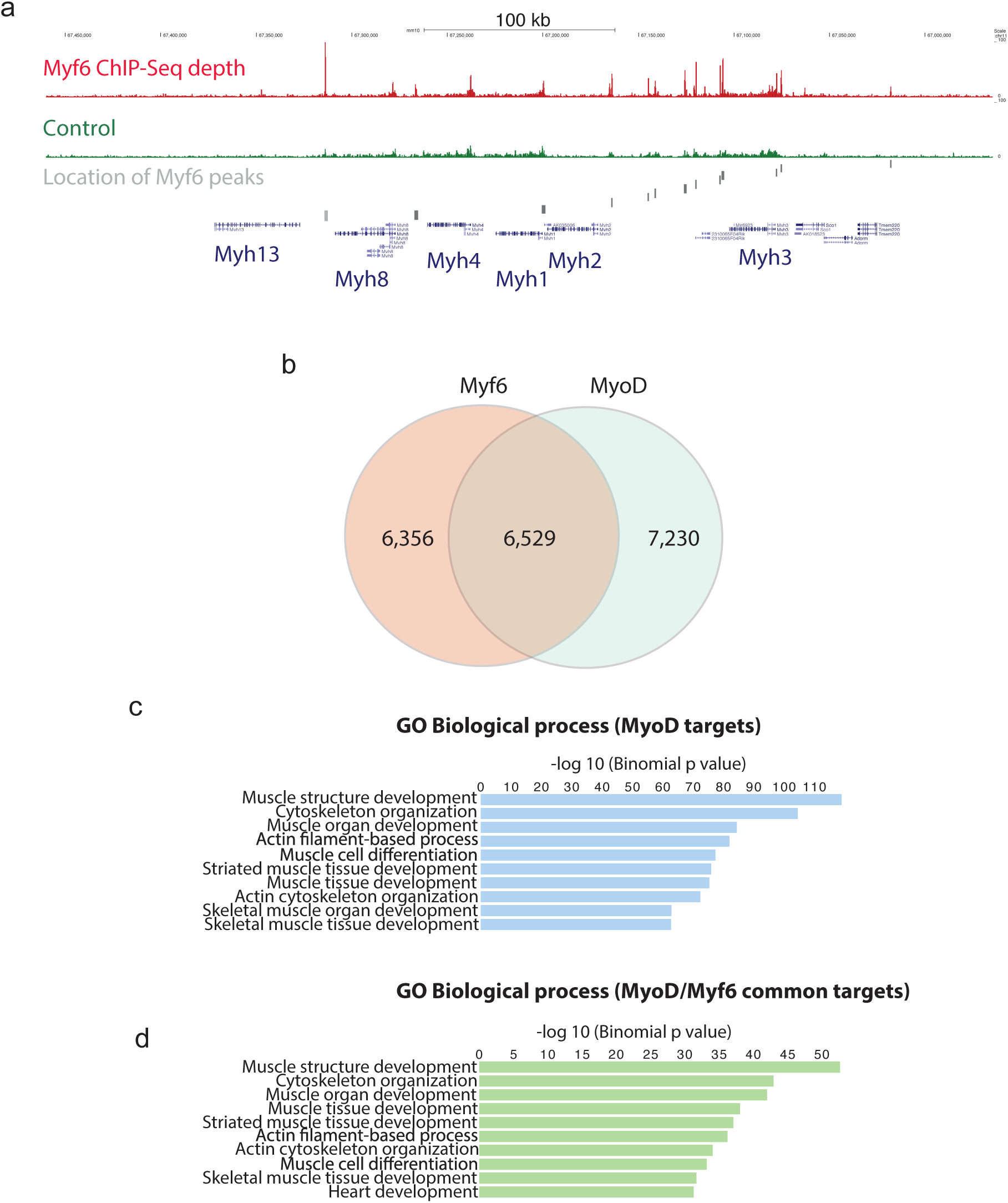
Myf6 and MyoD Share a Vast Core Myogenic Genetic Network. **(a)** Enrichment of Myf6 ChIP-Seq signal on the myosin heavy chain gene cluster and location of Myf6 peaks. **(b)** Venn diagram showing genome-wide binding sites for MyoD and Myf6 and their overlap in differentiated muscle cells. The overlap is defined as any peaks that has any overlap between the two datasets **(c)** Gene Ontology (GO analysis) using Genomic Regions Enrichment of Annotations Tool (GREAT) (McLean et al., 2010) of genes associated with MyoD ChIP-Seq peaks in primary myotubes (Soleimani et al., 2012a). **(d)** Similar analysis as in “c” for genes associated with MyoD/Myf6 common peaks. The association rule is based on single nearest gene within 300kb of the peaks. Within that window the number of peaks associated with more than one gene is minimal for ChIP-Seq datasets.

